# Spatial Transcriptomics Reveals the Temporal Architecture of the Seminiferous Epithelial Cycle and Precise Sertoli-Germ Synchronization

**DOI:** 10.1101/2024.10.28.620681

**Authors:** Arun Chakravorty, Benjamin D. Simons, Shosei Yoshida, Long Cai

## Abstract

Spermatogenesis is characterized by the seminiferous epithelial cycle, a periodic pattern of germ cell differentiation with a wave-like progression along the length of seminiferous tubules. While key signaling and metabolic components of the cycle are known, the transcriptional changes across the cycle and the correlations between germ cell and somatic lineages remain undefined. Here, we use spatial transcriptomics via RNA SeqFISH+ to profile 2,638 genes in 216,090 cells in mouse testis and identify a periodic transcriptional pattern across tubules that precisely recapitulates the seminiferous epithelial cycle, enabling us to map cells to specific timepoints along the developmental cycle. Analyzing gene expression in somatic cells reveals that Sertoli cells exhibit a cyclic transcriptional profile closely synchronized with germ cell development while other somatic cells do not demonstrate such synchronization. Remarkably, in mouse testis with drug-induced ablation of germ cells, Sertoli cells independently maintain their cyclic transcriptional dynamics. By analyzing expression data, we identify an innate retinoic acid cycle, a network of transcription factors with cyclic activation, and signaling from germ cells that could interact with this network. Together, this work leverages spatial geometries for mapping the temporal dynamics and reveals a regulatory mechanism in spermatogenesis where Sertoli cells oscillate and coordinate with the cyclical progression of germ cell development.

**Graphical Abstract:** 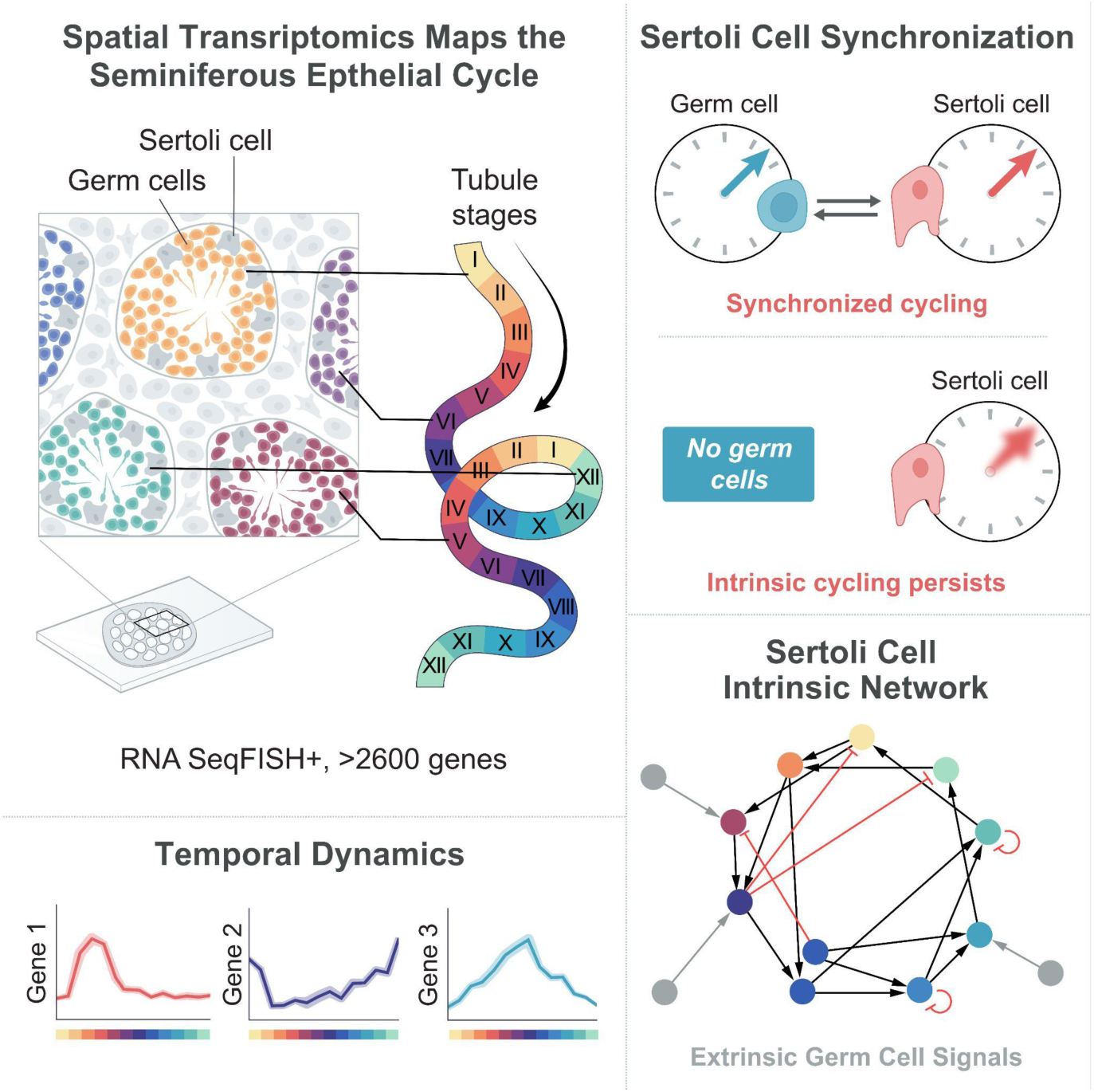

## Introduction

Spermatogenesis is the highly orchestrated, complex process that involves the transition of diploid spermatogonia into haploid spermatozoa, the male gametes essential for sexual reproduction. This intricately regulated process occurs within the seminiferous tubules of the testis and involves multiple stages of cellular differentiation, meiosis, and morphological remodeling. Each stage is marked by a unique set of cellular events and gene expression profiles, the regulation of which is still not fully understood.

Recent advancements in single cell RNA sequencing technologies have unveiled the transcriptional landscapes during spermatogenesis, illuminating marker genes and transient states pivotal to the process^1–4^. Yet, scRNA-seq is not without its limitations with tissue dissociation often resulting in a selective loss of cell types, particularly somatic cells in the testes. Moreover, while scRNA-seq excels in capturing transcriptional states, it surrenders information on the spatial cellular organization and relationships within the seminiferous tubules—a critical component when examining spermatogenesis. This spatial dimension, characterized by intricate geometries along two principal axes, plays a pivotal role in guiding cell-cell interactions, providing positional cues, and ensuring the systematic progression of differentiation.

Within the context of the seminiferous tubules, the phases of germ cell differentiation are separated along two principal axes. The first, the seminiferous epithelial cycle, exhibits a laminar differentiation geometry; in a cross section of seminiferous tubule, stem cells and spermatogonia are positioned along the basement membrane and as these cells differentiate, they move towards the central lumen, transitioning through various stages of development ^5,6^. This arrangement mirrors laminar differentiation geometry found in other tissues such as the skin interfollicular epithelium, the dentate gyrus of the hippocampus, and the retina^7–9^. The key stages of germ cell differentiation, including stem cell commitment, meiosis initiation, translocation from the basal to adluminal compartment, and spermiation, are coordinated by periodic signaling through retinoic acid^10^. In mice, this entire cycle takes ∼8.6 days and is divided into twelve stages, I to XII, and is defined by combinations of different cell types of maturing germ cells^5^.

A second principal axis of differentiation exists along the length of a seminiferous tubule, known as the spermatogenic wave. Here, the twelve stages of the periodic seminiferous epithelial cycle are organized in a sequential and periodic manner along the length of a seminiferous tubule, forming a wave-like pattern ^11,12^. This organization ensures the staggered and continuous production of mature sperm, underscoring the importance of spatial coordination in this intricate process. While the spermatogenic wave in rodents appears as successive stages of the epithelial cycle along the seminiferous tubule, the process is organized in more complex configurations in other species^13^. Nevertheless, the existence of the seminiferous epithelial cycle and the spermatogenic wave has been demonstrated in all amniote species examined, including humans^14^. Therefore, these spatial geometries are key conserved elements of spermatogenic regulation and play an important role in male fertility.

A recent endeavor in the realm of spatial transcriptomics captured this intricate spatial organization of spermatogenesis using Slide-seq, offering a resolution of 10uM^15,16^. While these studies shed light on the cellular compositions of spermatogonial microenvironments and stage-dependent gene expression patterns, each bead would capture mRNA from 2-3 adjacent cells, and often different cell types, limiting the ability to truly analyze single cell interactions. Moreover, these studies also did not capture the precise temporal architecture of the seminiferous epithelial cycle.

Here, we apply seqFISH+^17–19^, a method that resolves mRNA transcripts at a single molecule resolution, to adult mouse testis to study spermatogenesis. Using entire testis cross-sections, we systematically identified 26 cell types across 216,090 cells with 2638 genes, comprising of genes across all major signaling pathways and 1300 transcription factors. Since each tubule cross section is a time point in the seminiferous epithelial cycle, by mapping decoded transcripts to each tubule and computing their temporal order we reconstructed the seminiferous epithelial cycle with precise population dynamics. At a high dimensional level the tubules present a circular transcriptional topology, an expected feature of periodic transcriptional patterns. An analysis of somatic cells reveals that Sertoli cells exhibit a cyclic transcriptional profile closely synchronized with that of germ cells along the seminiferous epithelial cycle. In contrast, other somatic cells like peritubular cells and Leydig cells do not demonstrate such synchronization. By selectively ablating all germ cells using busulfan, a DNA alkylating agent, we find that Sertoli cells can autonomously maintain this cyclic transcriptional profile, suggesting an innate cyclic program tied to spermatogenesis. However, without germ cell communication we see varying degrees of gene dephasing and transcriptional downregulation predominantly involving PI3K-Akt, MAPK and Notch signaling. Finally, we identify an innate retinoic acid cycle in Sertoli cells as well as a network of transcription factors showing cyclic and directed activation suggesting a molecular mechanism driving the autonomous periodic transcriptional patterns that persist in Sertoli cells and identify how germ cells could reinforce the innate Sertoli cycle through signaling pathways.

Together, our results provide a detailed view of spermatogenesis at a single cell level and leverage the spatial geometries of tubules to obtain temporal dynamics along the seminiferous epithelial cycle. This approach yields a model that captures the intricate geometric and temporal patterns inherent to the seminiferous epithelial cycle. Furthermore, our finding that Sertoli cells display an autonomous cyclic transcriptional profile synchronized with germ cell development suggests a mechanism driving the coordinated timing of spermatogenesis.

## Results

### SeqFISH+ identifies 26 major cell types in the adult mouse testis at a single cell resolution

To study gene expression across the different cell types in the mouse testis, particularly within major signaling pathways, we identified 2638 genes encompassing a broad range of signaling pathways. This list included 1300 transcription factors and numerous cell-type specific markers (Methods, Supplementary Table 1). SeqFISH+ was performed on testis cross sections from two 8-week-old adult C57BL/6 mice. In total, we decoded >120m transcripts across 216,090 cells, at an average of 494.8 counts/cell. Furthermore, the data showed high reproducibility between batches (r = 0.98) (Figure S1A) and a strong correlation with bulk RNA seq data of adult mouse testes (r = 0.82) (Figure S1B). Across both batches, the mean false positive rate was 0.60% (Fig. S1C,D,E).

For cell segmentation we used concanavalin A(conA), a lectin that selectively binds to cell membrane glycoproteins a-mannopyranosyl and a-glucopyranosyl, thereby enabling accurate identification of cell membranes (Methods). To obtain cell segmentation masks we trained a Cellpose 2.0 model to use both conA and the DAPI stains to obtain precise cell segmentation masks ^20,21^ (Fig. S2C and Methods). Transcripts were then assigned to each cell based on the cell-masks.

After performing preprocessing and clustering steps (Methods), a UMAP projection revealed a continuum of germ cell populations alongside distinct clusters representing somatic cell types (Figure 1B, S1H). Using previously identified cell-type markers, we successfully identified all major germ cell populations covering the full developmental spectrum, in addition to all major somatic cell populations (Figure 1D). Notably, the extensive dataset allowed us to identify subcategories within undifferentiated spermatogonial stem cell populations (SSCs) (Fig. 1D, S3). These included SSCs primed for differentiation, as indicated by the expression of genes including *Neurog3, Sox3, and Sall1*. We also identified SSCs primed for self-renewal, marked by *Gfra1* expression. Additionally, we identified SSCs with an intermediate expression profile, suggesting a transitional state between renewal-primed and differentiation-primed states, a finding that aligns with our recent study^22^. Importantly, our dataset uniquely captured a broad range of somatic cell types, including Sertoli, Leydig, peritubular, macrophages, endothelial, and perivascular cells, which are often underrepresented in scRNA-seq datasets. By mapping cell-type identities back to their spatial coordinates, we observed a strong concordance between cellular identity and expected anatomical location (Figure 1C, S2B). For instance, spermatogonia including SSCs were consistently localized at the tubule border, while spermatocytes were found in the adjacent layer. Round spermatids were found in the following inner layer, and elongated spermatids showed closest localization to the tubular lumen. Alongside the stem cells and spermatogonia, Sertoli cells were interspersed along the basement membrane, with peritubular cells delineating the tubule border. Finally, in the interstitial spaces we found somatic cells including macrophages, Leydig, perivascular, and endothelial cells. In rare instances we also found examples of T-cells, all restricted to the interstitial spaces. Thus, our dataset not only enabled the identification of all major cell types but also facilitated the delineation of previously unappreciated subtypes, all while mapping these cells to their precise anatomical locations.

**Figure 1:**
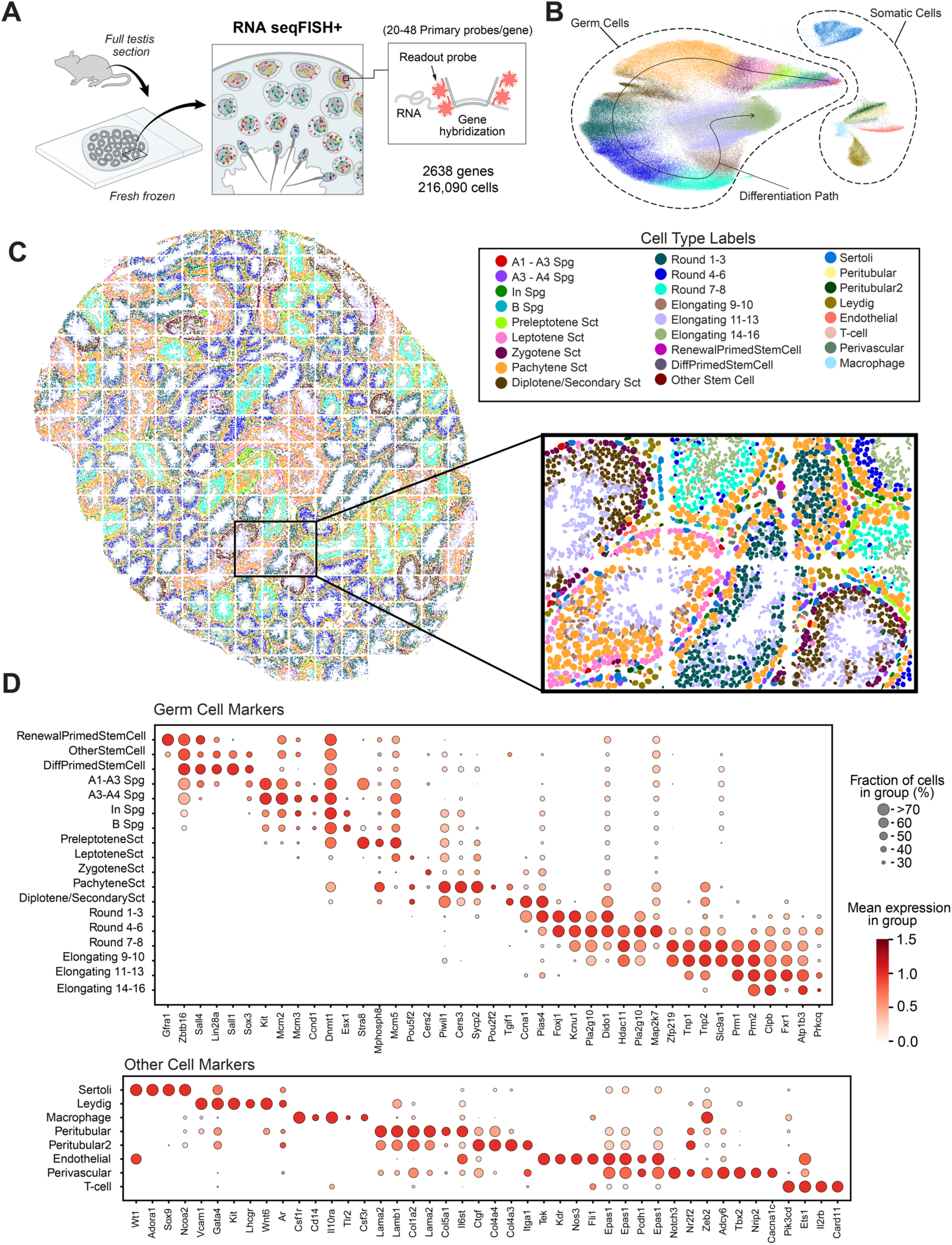
SeqFISH+ resolves 26 major cell types at single-cell resolution in wildtype mouse. **(A)** Experimental workflow: Fresh frozen cross-sections of 8-week-old adult mouse testes were imaged using seqFISH+ for a panel of 2638 genes. Primary probes hybridize directly to the complementary target RNA, and readout probes bind to the primary probe, resolving each RNA at a single-molecule resolution. Over 216,090 cells were identified. **(B)** Uniform Manifold Approximation and Projection (UMAP) visualization of integrated seqFISH+ datasets. Cluster colors are consistent with panel C. Germ cells form a continuum reflecting their differentiation trajectory, while somatic cells cluster separately. **(C)** A spatial map from a full testes cross section showing the locations of 26 different cell types identified by transcriptional profile. Stem cells and other spermatogonia were found along the basement membrane of the seminiferous tubules, while spermatocytes and spermatids were found to be organized laminarly towards the tubular lumen. Sertoli cells were found spaced out along the basement membrane of each tubule while Leydig cells, perivascular, and endothelial cells were found in interstitial spaces. Finally, two types of peritubular cells were identified forming the border of each tubule cross section. A detailed visual of each cell type is presented in Figure S2. **(D)** Marker genes for each of the identified cell types. Upper panel focuses on germ cell markers while the lower panel focuses on markers for the somatic cell populations. *See Figure S1 and Figure S2 for additional data*.

When viewing all cell types across the full testes cross section, it became evident that each individual tubule possesses a distinct cellular composition (Figure 1C). This variability aligns well with the concept that each tubule-cross section represents a unique stage of development thereby echoing the established dynamics of the seminiferous epithelial cycle. For example, tubules that were identified to be at Stage VII-VIII based on expression of *Stra8* and DAPI nuclear morphology, showed presence of early differentiating spermatogonia, preleptotene spermatocytes, pachytene spermatocytes, and elongating spermatids. In contrast, tubules at Stage I-III showed the presence of early round spermatids, a hallmark of these stages. Thus, each tubule manifests a characteristic stage of differentiation that is consistent with the seminiferous epithelial cycle.

### A tubule-level analysis reveals the temporal architecture of seminiferous epithelial cycle

From our initial observation that each tubule cross-section has a distinct cellular composition, we deduced that individual tubule cross sections represent discrete time points along the seminiferous epithelial cycle. To explore this temporal architecture further, we aggregated transcript counts for each tubule to analyze our data at a tubule level (Fig. 2B). Post-normalization and scaling (Methods), we found that the tubules organized into a circular transcriptional topology in PC space (Fig. 2D). This topology is reminiscent of other cyclic biological processes such as the cell cycle and circadian rhythm.

**Figure 2:**
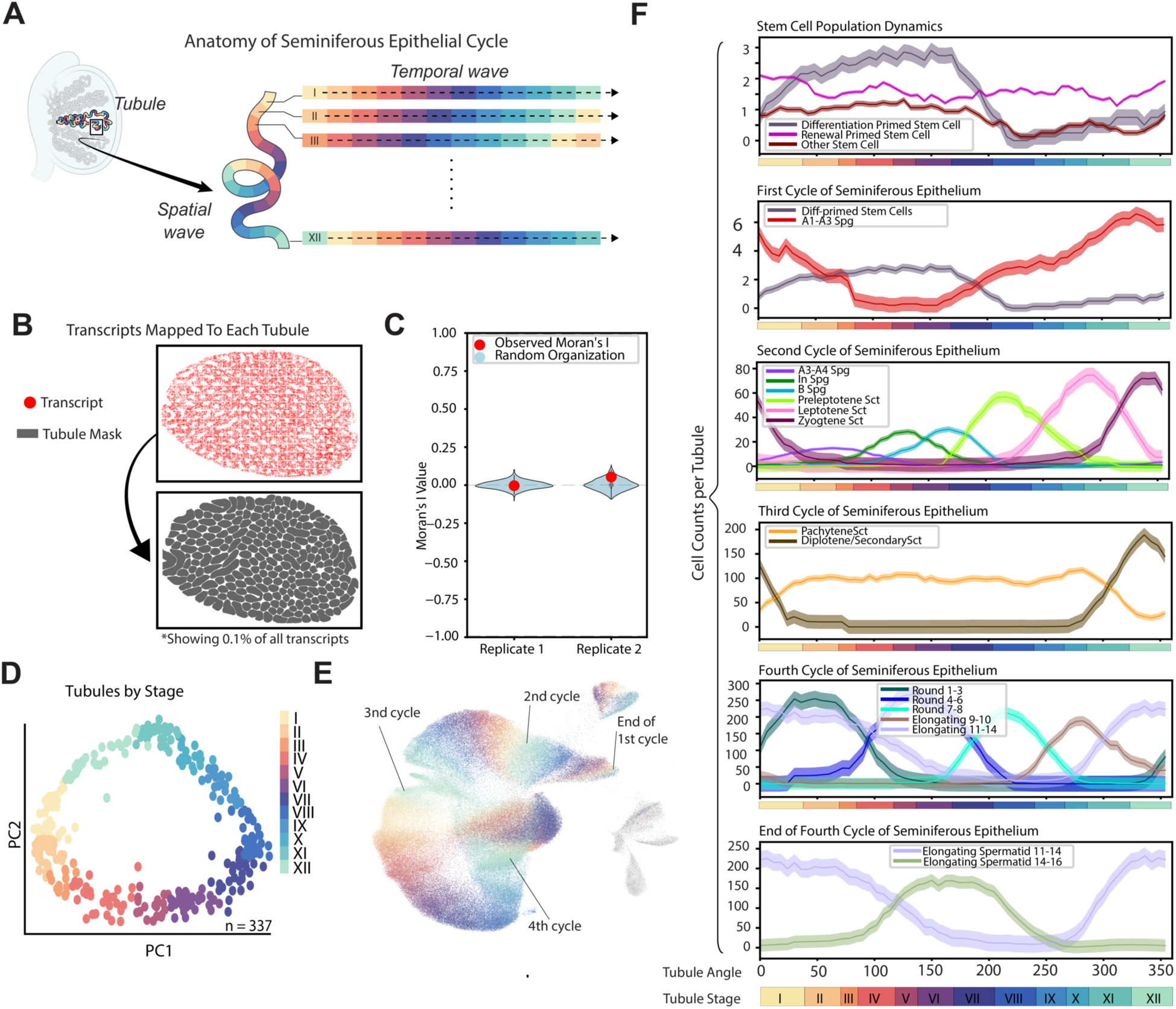
A tubule-level analysis reveals the temporal architecture of the seminiferous epithelial cycle. **(A)** Visualization of the seminiferous epithelial cycle. Along each seminiferous tubule, stages of the seminiferous epithelial cycles are represented periodically (I - XII). As time passes, each cross section of the tubule progresses through the stages of seminiferous epithelial cycle. **(B)** Transcripts per tubule were mapped based on tubule masks and the resulting tubule x gene matrix was analyzed. **(C)** A modified Moran’s I for cyclic parameters, *I_cyclic,_* was used to evaluate whether Tubule Angle was spatially organized across testis cross sections. For both replicates, the tubule organization was found to be within random range, with an *I_cyclic_* =-0.004 and 0.054 for each replicate respectively. **(D)** Principal component analysis showed a circular transcriptional profile in PC1 and PC2, which was found to be representative of the cyclic process of the seminiferous epithelial cycle. **(E)** Approximately 4 cycles of the seminiferous epithelial cycle are required for stem cells to mature into spermatids. A UMAP representation of the full dataset shows the 4 cycles of the seminiferous epithelial cycle, with cells colored by their corresponding tubule stage. **(F)** Cell counts per tubule, mean +/- SEM, for each cell type along each cycle of the seminiferous epithelium. Four complete cycles are required for stem cells to mature into spermatids. *See Figure S4, Figure S5, and Figure S6 for additional data*.

To determine whether the circular transcriptional profile represents the entire seminiferous epithelial cycle, in which tubules cyclically transition from Stage I through Stage XII (Fig. 2A), we investigated the population dynamics of cells along this circular transcriptional profile. We first computed an ordering of the tubules using scPrisma, a package designed to find the order of cells along cyclic processes using topological priors^23^ (Fig. S5B, S5C). These orderings were then converted to angles uniformly from 0 degrees to 360 degrees for a more universal metric, called ‘Tubule Angle’ (Methods) (Fig. S5B, S5D). Subsequently, we plotted the known cell-type populations of each tubule, derived from our spatial transcriptomics dataset, as a function of the Tubule Angle (Fig. 2F). We reasoned that if the circular transcriptional profile represents the seminiferous epithelial cycle, we would see sequential and predictable changes in the prevalence of different cell types corresponding to each stage of the cycle. For instance, we would expect a decline in spermatogonia populations, due to differentiation, to coincide with an increase in spermatocyte numbers. Similarly, tubules with round spermatids would be followed by those enriched for elongating spermatids.

Indeed, our data corroborated these expectations precisely. A consistent germ cell ordering— from A1-A3 spermatogonia to elongated spermatid 14-16—was evident along the tubule sequence (Fig. 2F). Importantly, each cell type displayed a correlated decrease in prevalence concomitant with a rise in its immediate derivative cell type. This pattern held true across all stages of cell differentiation and maturation, demonstrating a continuous, stepwise progression along the tubule ordering.

Having established that the order of tubules in PC space represents the seminiferous epithelial cycle, we next sought to align these with classical stages, I - XII. First, we explored whether our testis cross-sections displayed any bias for specific tubule stages by quantitatively measuring spatial autocorrelation using a modified Moran’s I for cyclic parameters (Fig. 2C, Fig. S4, Methods). Consistent with histological observations ^11,24^, we found that tubule stages are not organized spatially in a testis cross section. Specifically, we found that the observed Tubule Angles were random with respect to the whole testis cross section, presenting robust evidence for the lack of global inter-tubular organization (Fig. 2C, Fig. S4). Thus, each tubule cross section represents a random discrete time point along the seminiferous epithelial cycle.

Given our large sample size of 337 tubule cross sections, we quantitatively validated that moving averages would reliably estimate the true temporal dynamics of the seminiferous epithelial cycle (Fig. S4F). Using known frequencies and timescales of each stage^11^ (Fig S5B, S5E), we mapped back our Tubule Angles to stages, validated by DAPI nuclear morphology (Fig. S6B), and found precise consistency between cell population dynamics and the known dynamics of seminiferous epithelial cycle (Fig. 2F). For instance, at stage VII we see the impact of retinoic acid causing: 1) the commitment to differentiation and the rise in *Kit+* A1-A3 spermatogonia; and 2) commitment to meiosis and the rise of *Stra8*+ preleptotene spermatocytes. As another example, at stage I we see the rise of populations of pachytene spermatocytes and round spermatids. Finally, our analysis suggests that four complete cycles of the seminiferous epithelium are required to transition from A1-A3 Spermatogonia to mature elongated spermatids, in perfect agreement with longstanding literature (Fig. 2E)^11,25^. Overall, all staging and cell type populations mapped precisely to known populations of the seminiferous epithelial cycle, and an overall schematic is shown in Figure S6A.

Notably, we were able to profile the population dynamics of SSCs across the different subtypes identified: renewal-primed (*Zbtb16*+, *Sox3*-, *Sall3*-, *Gfra1*+), differentiation-primed (*Zbtb16*+, *Sox3*+, *Sall3*+, *Gfra1*-), and the transitional/other stem cell populations (*Zbtb16*+, *Sox3*+, *Sall3*+, *Gfra1*+) (Fig. 2F, Fig S3A-H). We found that differentiation-primed SSCs increased from stage IX to VII but sharply declined at stage VII—coinciding with retinoic acid presence—and were replaced by *Kit*+ A1-A3 spermatogonia (Fig. 2F). Renewal-primed SSCs remained stable at ∼2 cell/tubule cross-section, unaffected by retinoic acid, with slight increases from stage XII to I.

Transitional/other SSCs, exhibiting both self-renewal and early differentiation markers, were relatively stable but also decreased at stage VII suggesting a potential impact of retinoic acid on the transition of stem cells between renewal and differentiation-primed states. These stage-dependencies show striking agreement with cell type distribution of SSCs defined by protein expression^22^, indicating the transcriptional states faithfully capture the phenotypes in these cells.

### Sertoli Cells display a dynamic transcriptional profile linked to the seminiferous epithelial cycle

Mammalian testes are characterized by a complex cellular landscape, consisting of a diverse repertoire of both germ cells and somatic cells. While the classical staging of the seminiferous epithelial cycle has primarily focused on the developmental stages of germ cells, our understanding of the transcriptional dynamics of somatic cells through the cycle remains poorly characterized.

When Sertoli cells were clustered independently of germ cells, we observed a circular transcriptional profile in the gene expression space, which strikingly resembled the cyclical nature observed in the seminiferous epithelial cycle. Furthermore, by mapping these Sertoli cells backs to their respective Tubule Angle — based on their physical location within individual tubules — we found a high degree of congruence between the circular manifold and the Tubule Angle (Fig. 3A). This shows that the transcriptional landscape of Sertoli cells is intricately linked to the stages of the seminiferous epithelial cycle (Fig. 3B), consistent with lines of existing evidence that have noted stage-dependent expression in Sertoli cells ^1,15,26–30^. The clear cyclical expression profile observed in our dataset was uniquely captured due to the unbiased sampling of Sertoli cells across all stages of the cycle. Unlike traditional single-cell approaches where the complex branched architecture of Sertoli cells makes them particularly vulnerable to damage during tissue dissociation, resulting in their significant underrepresentation in single-cell datasets^1,4^, our spatial method preserves their architecture and enables comprehensive profiling spanning thousands of cells in each replicate.

**Figure 3:**
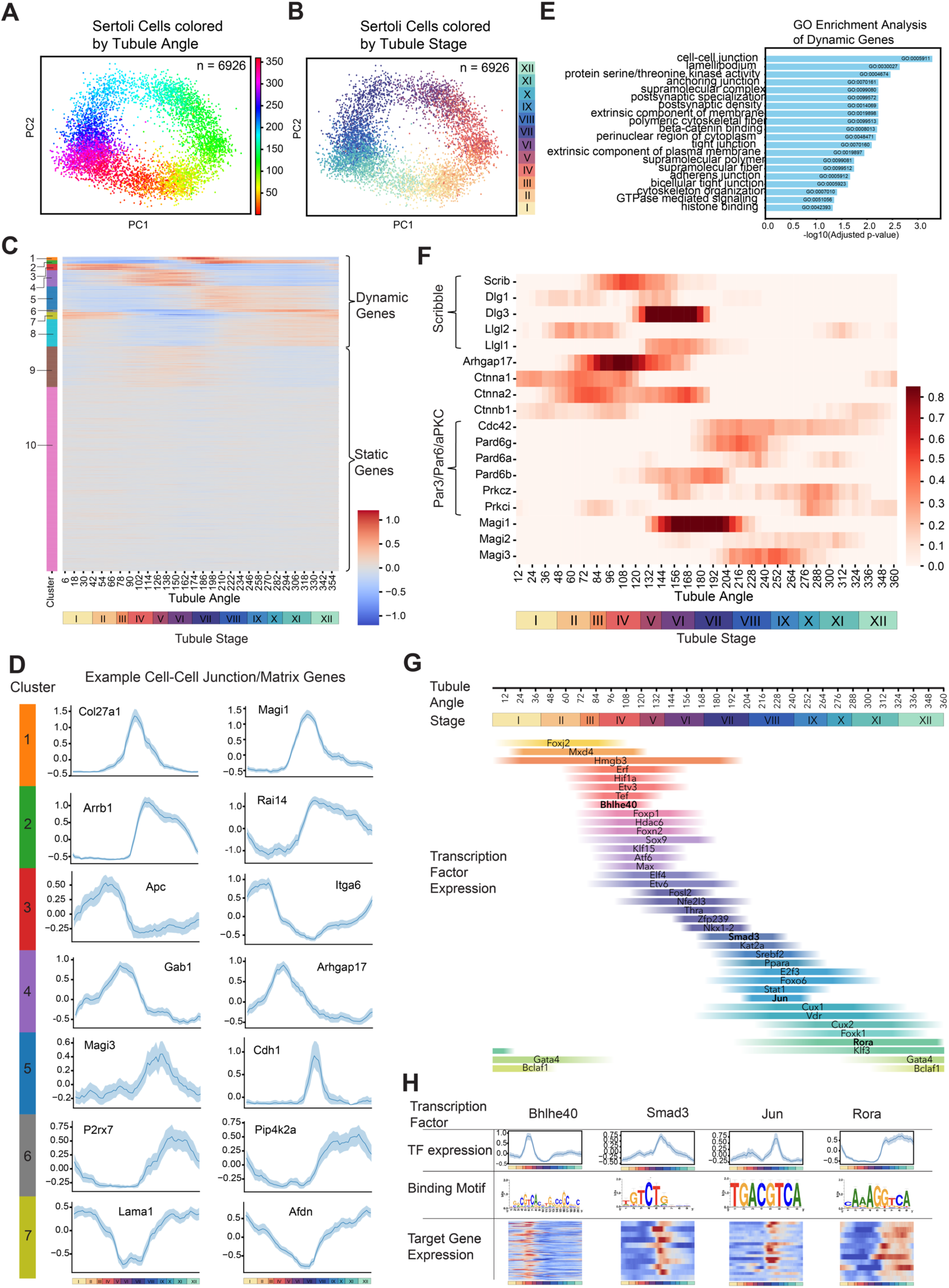
Sertoli Cells display a cyclic transcriptional profile synchronized with the seminiferous epithelial cycle. **(A, B)** Sertoli cells were isolated and analyzed. In PC space, the transcriptional profile showed a circular topology. When colored by the Tubule Angle and Tubule stage (stages being a discretized version of Tubule Angle) from which the Sertoli cells originated, the cyclic transcriptional topology appears to be synchronized with the seminiferous epithelial cycle. **(C)** All 2,638 genes expressed in Sertoli cells were clustered based on their expression profile along Tubule Angle/Tubule Stages. Many clusters (1-7) showed specific phase-dependent expression patterns, while other clusters showed no phase-dependent patterns. **(D)** Example expression profiles of genes related to cell-cell junctions or extracellular matrix components in Sertoli cells from clusters 1-7 in subpanel C. 95% confidence intervals were evaluated by randomizing time points and determining the interval for noise. **(E)** Gene ontology analysis for the dynamic genes in clusters 1-7 showed enrichment in terms associated with junction complexes and extracellular matrix formation, such as cell-cell junction, anchoring junction, tight junction, bicellular tight junction, and extracellular membrane components, among others. **(F)** Heatmap showing the cyclic expression patterns of genes related to the Scribble and Par3/Par6/aPKC polarity complexes in Sertoli cells across the seminiferous epithelial cycle. **(G)** The role of transcription factors on expression of target genes was evaluated using the Scenic pipeline^41^. Transcription factors with target genes with clear phase dependent expression were identified and summarized. **(H)** Examples of 4 regulons for transcription factors *Bhlhe40, Smad3, Jun*, and *Rora*. Each transcription factor and its respective target genes displayed phase-dependent expression patterns. The specific binding motif found to be enriched for the target genes specific to each transcription factor is also shown.

To further characterize this dynamic expression landscape of Sertoli cells, we explored the gene expression profile as a function of Tubule Angle/stage of the seminiferous epithelial cycle. Among the 2638 genes profiled in our study, over 700 genes displayed dynamics visibly tied to the seminiferous epithelial cycle (Fig. 3C, Fig. 4G). Importantly, many of these genes clustered into groups, suggesting the involvement of broad signaling mechanisms in these expression changes.

**Figure 4:**
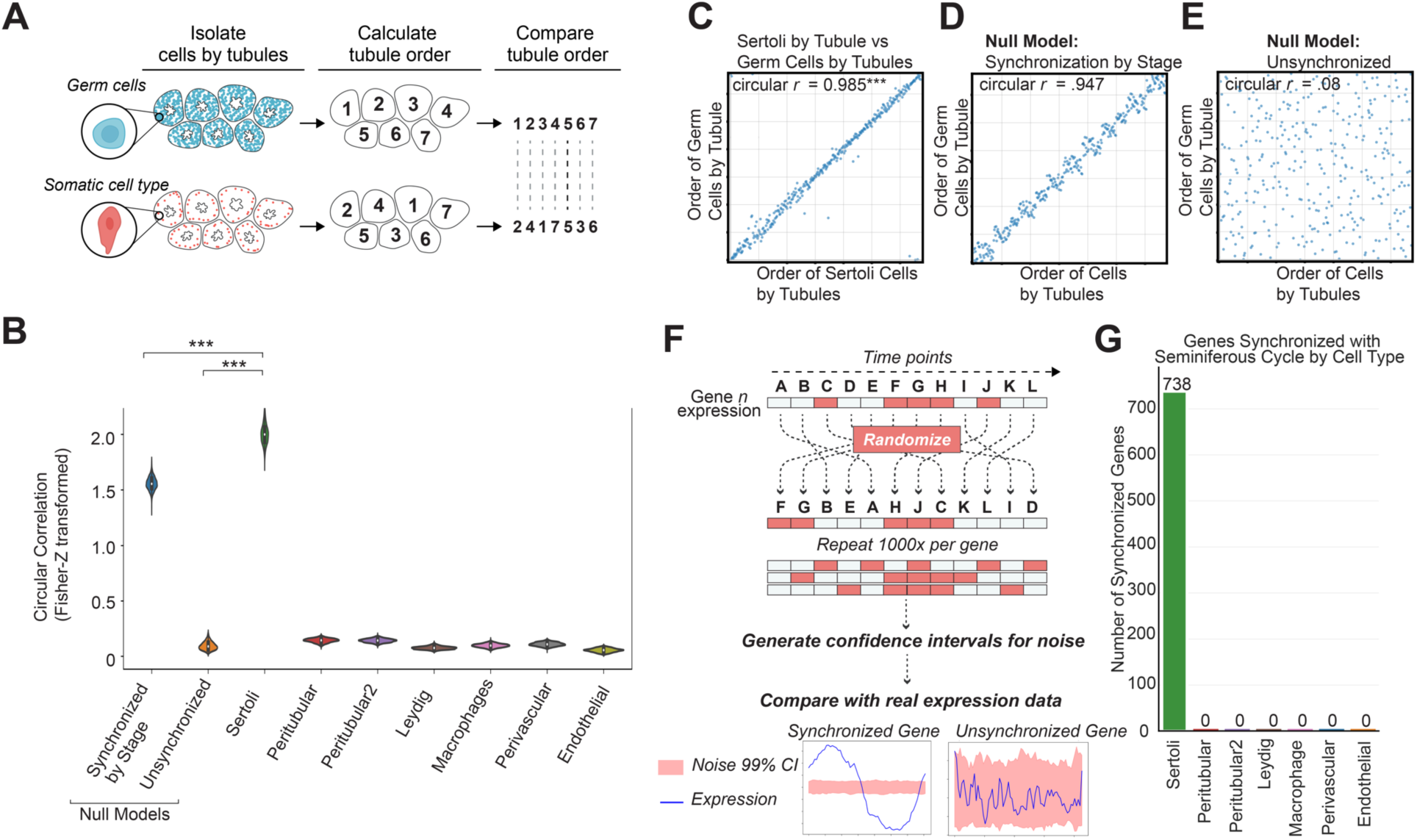
Sertoli cells are the only somatic cells synchronized with the seminiferous epithelial cycle. **(A)** Each somatic cell type was isolated and grouped by the tubule they were in or closest to. Similarly, germ cells were grouped by tubule. Using the scPrisma pipeline (Methods), the order of the tubules for the germ cell groupings and the somatic cell groupings were independently calculated and compared. The degree of correlation between the independent orderings quantifies the degree of synchronization between the somatic cell type and the germ cells at a topographic level. These independent orderings were computed one thousand times for each cell type. **(B)** Pearson circular correlation was computed between the independent orderings of each somatic cell type and germ cell groupings by tubule. Only Sertoli cells displayed a high correlation, exceeding the correlation for the null models expected for Unsynchronized and Synchronized by Stage phenotypes (p = 0.000, p = 0.000 respectively). On the other hand, all other somatic cell types, peritubular, peritubular2, Leydig, macrophage, perivascular, endothelial displayed a correlation that was not statistically different from the Null model for Unsynchronized behavior (p= 1.000, 0.998, 1.000, 1.000, 1.000, 1.000, respectively). **(C)** Example iteration of independent ordering of Sertoli cell groupings compared with independent ordering of germ cell groupings. **(D)** Example iteration of germ cell orderings compared with null model for synchronization by stage. **(E)** Example iteration of germ cell orderings compared with Unsynchronized null model. **(F)** To evaluate the synchrony of specific genes, the 99% confidence interval of noise was computed by scrambling the time points (Tubule Angle) randomly. The true expression profiles were then compared to the profiles for noise to determine which genes displayed expression that were phase-dependent, or synchronized with the seminiferous epithelial cycle. An example of synchronized behavior and unsynchronized expression profiles compared to noise is shown. **(G)** Only Sertoli cells showed gene expression profiles outside the expected range of noise. 738 genes showed expression profiles characterized as synchronized to the seminiferous epithelial cycle. For all other somatic cell types no genes were found to be synchronized to the seminiferous epithelial cycle.

Our Gene Ontology (GO) analysis revealed functional relevance to these dynamically expressed genes, implicating various types of cellular junctions (e.g., tight junction, cell-cell junction, anchoring junction) and morphological changes (e.g., lamellipodium, extrinsic component of membrane, cytoskeletal organization) and phosphorylation-based events (Fig. 3D,E). The dynamic expression of genes related to cellular junctions and extracellular matrix modifications reinforces the active role Sertoli cells play in supporting and guiding the spermatogenic process ^31–33^.

Among genes related to junction activity, we found orthogonal expression in two pathways, Scribble and Par3/Par6/aPKC, that are reciprocally inhibitory and related to establishment of cell polarity. The Scribble polarity complex, a major basolateral polarity regulator composed of Scribble(*Scrib),* Discs Large(*Dlg1, Dlg3*), and Lethal Giant Larvae(*Llgl1*, *Lgl2*) were found to be expressed between stages II - VII (Fig. 3F). In contrast, expression of Par3/Par6/aPKC complex, known for conserved set of apical polarity factors including *Pard6g*, *Pard6a, and Pard6b,* part of the Par6 complex, and *Prkcz*, an aPKC member, were found to be expressed from stages VII - XI (Fig. 3F). This alternating pattern suggests a dynamic remodeling of the cellular junctions forming the blood-testis barrier (BTB). The expression of the Scribble complex proteins at earlier stages(II-VII) suggests a role in maintaining the basolateral domain, consistent with its role in epithelial cells ^34,35^. The subsequent expression of Par complex proteins from Stage VII coincides with the known period of BTB restructuring ^31^, likely driving the remodeling of tight junctions apically as seen in other systems ^34,36^. This temporal sequence would facilitate the migration of preleptotene/leptotene spermatocytes from the basal to adluminal compartment around stage IX, while preserving overall barrier integrity. Supporting this model, *Cdc42*, essential for activating the Par complex ^34^, and Magi proteins (*Magi1, Magi3*), crucial for proper localization of Par complexes apically^37^, were also expressed from Stage VII - XI(Fig. 3F). Following spermatocyte migration into the adluminal compartment, tight junctions are re-established at the basolateral domain driven at least partly by the Scribble pathway. In addition, *Arhgap17* (also *Rich1*), a Cdc42-specific RhoGAP^38,39^, is specifically expressed from stages III - VI (Fig. 3F), likely controlling the activity of Cdc42, maintaining junctions laterally^40^. This process creates a “treadmill-like” movement of tight junctions that allow migration of spermatocytes while maintaining BTB integrity.

To characterize the gene regulatory networks of Sertoli cells we identified transcription factors(TFs) and target genes that displayed enrichment of the respective binding motif and similar expression profiles^41^ (Fig. 3G, 3H). These TFs included marker genes such as *Sox9, Gata4, Foxp1,* among others. While some of these genes have been noted to display stage specific expression ^42,26^, the precise role of these TFs is yet to be explored systematically.

### Sertoli cells are the only somatic cells precisely synchronized with the seminiferous epithelial cycle

While Sertoli cells displayed a cyclic transcriptional profile that was clearly linked to the seminiferous epithelial cycle, we sought to uncover the degree of synchronization and whether other somatic cells shared similar synchronization. To quantify the correlation, we grouped somatic cell types and germ cells according to their associated or nearest tubules and then independently calculated and compared the tubule ordering for both groupings using scPrisma^23^ (Fig. 4A). The degree of correlation between these independent orderings quantifies the synchronization between each somatic cell type and germ cells at a broad topographic level.

Our results showed that Sertoli cells were the only somatic cells that showed a high correlation between tubule orderings (Fig. 4B). Indeed, Sertoli cells showed a level of synchronization significantly greater than our null model, which was synchronized at a stage level. This suggests that Sertoli cells are precisely synchronized to the dynamics of the seminiferous epithelial cycle along a continuum, rather than limited to state transitions. In contrast, all other somatic cells including peritubular, Leydig, macrophage, perivascular, and endothelial cells, showed correlations comparable to our null model for unsynchronized behavior (Fig. 4B, Fig. 4E).

We further quantified whether any genes in all somatic cell types displayed dynamics that were synchronized with the seminiferous epithelial cycle by comparing their expression profiles to our noise estimates, obtained from scrambling the time points (Fig. 4F). Consistent with our previous findings, only Sertoli cells had genes that were synchronized with the seminiferous epithelial cycle while no genes were found to be synchronized in all other somatic cell types (Fig. 4G).

### Sertoli cells autonomously maintain a cyclic transcriptional profile without the presence of germ cells

Having established the unique synchronization of Sertoli cells with the seminiferous epithelial cycle we next sought to address a fundamental question: are Sertoli cells the drivers of this cyclic program, or are they merely responding to signals from the germ cells? While traditionally it has been thought that Sertoli cells are guided by germ cells and simply provide support, other results have brought this into question, suggesting a potential innate cyclical program^43^. Thus, we devised an experiment to observe Sertoli cell behavior in the absence of germ cells. We hypothesized that if Sertoli cells maintain their cyclic transcriptional profile without germ cells, it would suggest an intrinsic program within Sertoli cells themselves.

Using busulfan, a DNA-alkylating agent commonly used for ablation and transplantation experiments, we ablated all differentiating germ cells in wildtype mice ^44,45^(Fig 5A). Importantly, since all Sertoli and other somatic cells are not dividing, this treatment does not affect them. Thirty days after treatment, the busulfan treated testes revealed a stark contrast compared to wild-type, displaying empty tubules while Sertoli and other somatic cells remained (Fig. 5B-C, Fig. S7). Remarkably, after plotting Sertoli cells grouped by tubule we saw a discernible cyclic topology in PC space (Fig. 5D). Furthermore, when these tubules were plotted along untreated Sertoli groupings, we saw that the busulfan treated Sertoli groupings occupied a cyclic transcriptional profile within the cyclic transcriptional profile of the untreated Sertoli groupings (Fig. 5E).

**Figure 5:**
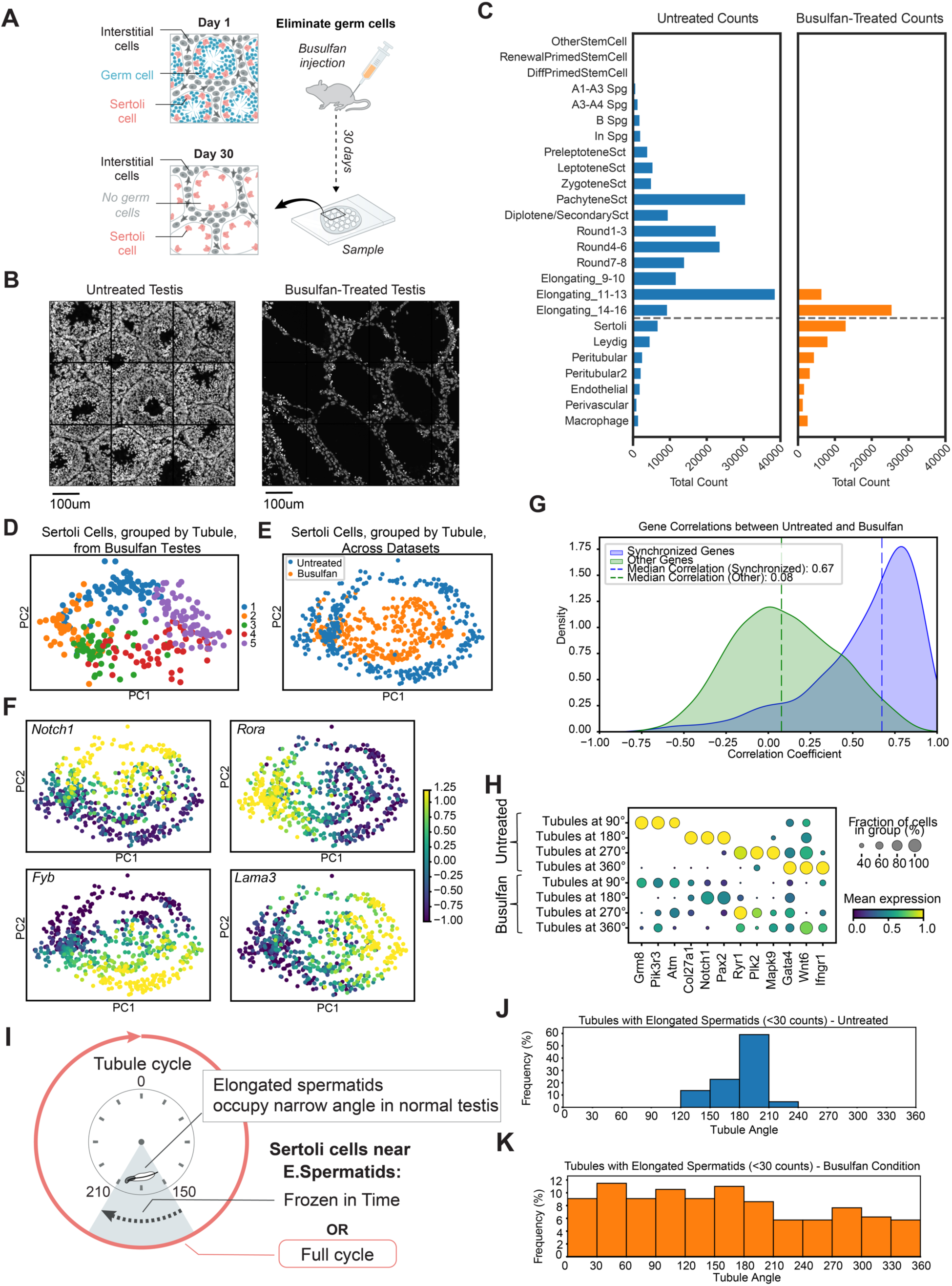
Sertoli cells independent of Germ cells maintain a cyclic transcriptional profile. **(A)** 5-6 week old wildtype mice were injected with busulfan at a concentration of 30 or 45mg/kg intraperitoneally. After 30 days, the testes were harvested and processed for seqFISH+ imaging. **(B)** DAPI images comparing untreated testis cross section vs a busulfan treated testis cross section. The busulfan treated testis cross sections showed tubules void of differentiating germ cells while somatic cells were still visible **(C)** Post-cell typing, the numbers of each cell type of untreated testis were compared with busulfan treated testis. Complete ablation of differentiating germ cells was observed with remnant elongated spermatids. Somatic cells were unaffected and retained similar population levels as untreated. **(D)** Sertoli cells in the busulfan treated testes were grouped by tubule and plotted in PC space using genes known to be dynamically expressed in the untreated condition. Each tubule was also scored by the expression of gene sets found to be expressed in a synchronized manner to the seminiferous epithelial cycle. Tubules were colored by the gene set that scored highest. Specifically gene set 1,2,3,4,5 consisted of genes from Cluster 1 & 2, 3, 4, 5 & 6, 7 from Figure 3C, respectively. **(E)** Sertoli cells averaged by tubule were merged from the untreated condition and busulfan condition, analyzed, and plotted in PC space. Sertoli cell groupings from the busulfan condition display a cyclic transcriptional profile within the confines of the cyclic profile of the untreated Sertoli cell groupings. **(F)** Expression of many genes showed expression in similar points in the manifolds of untreated and busulfan treated Sertoli cell groupings. Shown are example genes, *Notch1, Rora, Fyb, Lama3*, which were expressed in orthogonal phases of the seminiferous epithelial cycle in untreated Sertoli cells. **(G)** The order of tubules was computed independently for Sertoli cells grouped by tubule for the busulfan condition and optimally aligned to the order of untreated Sertoli groupings based on gene expression profiles. The circular Pearson correlation was then computed for each gene. Genes synchronized to the seminiferous epithelial cycle showed a high median correlation (r = 0.67) between the busulfan condition and the untreated condition, while genes identified as unsynchronized to the seminiferous epithelial cycle showed a low median correlation (r = 0.08), providing strong evidence that the cyclic transcriptional profile of Sertoli cells in the busulfan condition is similar to that of the cyclic profile of Sertoli cells in untreated tubules. **(H)** Gene expression patterns across distinct Tubule Angle +/- 2 degrees in Sertoli cells in the untreated condition versus busulfan condition. Across many genes a level of dephasing was noticeable. While Sertoli cells in untreated animals at 90, 180, 270, and 360 display distinct expression profiles, Sertoli cells from the busulfan condition display mixed expression profiles signifying a level of dephasing and blending between previously orthogonal expression profiles. **(I)** Graphical illustration for examining if Sertoli cells in the busulfan condition are frozen in time or continue to cycle. Elongated spermatids exist in untreated testis in Stage VIII tubules, roughly between Tubule Angle 150 - 210. In the busulfan condition, Sertoli cells in tubules with elongated spermatids were inspected. If Sertoli cells were frozen in time, we would expect those Sertoli cells to only match the Tubule Angle where elongated spermatids are typically found. If Sertoli cells continue to cycle after spermatids have matured completely and all other differentiating germ cells are absent, we would expect to see all Tubule Angles represented. **(J-K)** Distribution of Tubule Angles represented by tubules containing elongated spermatids, defined as elongating spermatids with fewer than 30 counts, in untreated and the busulfan condition. As expected, elongated spermatids in untreated are only seen in predominantly 180 - 210 degrees, coinciding roughly with Stage VIII and the completion of the spermatogenesis and prior to the release of elongated spermatids at the end of Stage VIII. In contrast, elongated spermatids are found at all Tubule Angles, suggesting the Sertoli cells continue to cycle even after spermatogenesis is complete and there are no longer any remaining differentiating germ cells. *See Figure S7 and Figure S8 for additional data*.

Across many genes the cyclic transcriptional profile of the Sertoli cells from the busulfan conditions displayed similar albeit less coherent patterns to the untreated condition suggesting that the gene programs still retained the same innate program (Fig. 5F). To quantify this, we computed the circular Pearson correlation for each gene between the two conditions (Fig. 5G). Genes that we had previously identified as synchronized to the seminiferous epithelial cycle showed a high correlation (median r = 0.67) between untreated and busulfan conditions. In contrast, genes that we had classified as unsynchronized showed a low correlation (median r = 0.08). This provided strong evidence that the cyclic transcriptional profile of Sertoli cells persists independently of germ cells.

The persistence of a cyclic transcriptional profile in Sertoli cells in the busulfan condition similar to that of Sertoli cells in the untreated conditions suggests that Sertoli cells are either continuing to cycle or are frozen in time. To investigate this we examined tubules in the busulfan condition containing remnant spermatids. If the Sertoli cells were indeed frozen in time, we would expect them to have gene expression patterns corresponding to the Sertoli cells that are present with elongated spermatids in the untreated condition (Fig 5I). For instance, in the untreated testis, elongated spermatids only occupy Tubule Angles between 180° - 210°, coinciding with the Stage VIII and the end of the 4th round of the seminiferous epithelial cycle (Fig 5J). However, in the busulfan condition, we found that tubules containing elongated spermatids were encompassing Sertoli cells occupying all Tubule Angles from 0° - 360° (Fig 5K), strongly suggesting Sertoli cells continue to transcriptionally cycle even after there are no longer any differentiating germ cells. This is also consistent with earlier findings where Sertoli cells in mice lacking differentiating germ cells show variable expression of stage-specific proteins, *galectin* and *tPa,* similar to those observed in normal testis, suggesting a cell-intrinsic cycle ^43^.

Our findings also revealed important differences in the Sertoli cells program without germ cells. When we examined gene expression patterns across specific tubule angles (Fig. 5H), we noticed a level of dephasing in the busulfan-treated condition. While Sertoli cells in untreated animals at 90°, 180°, 270°, and 360° tubule angles displayed distinct expression profiles, Sertoli cells from busulfan-treated testes showed more mixed expression profiles. This blending of previously orthogonal expression patterns suggests that while the overall cyclic program is maintained, the fine-tuning and precise synchronization of gene expression relies on the presence of germ cells.

### Sertoli cells retain intratubular synchronization in the absence of germ cells but gene expression levels dephase and specific signaling pathways downregulate

Since our previous analysis focused at a tubule level, the dephasing observed at a gene level brings two possible scenarios: 1) Sertoli cells of the same tubule were *desynchronized* causing the mixed expression profile of the tubule, 2) Sertoli cells at a single cell level develop mixed expression profiles due to *dephasing* of the cyclic expression profile (Fig. 6A). To answer this question, we analyzed the distance of the expression profiles of single cells within tubules vs across tubules. We expected that if case 1) were true, distance between cells within tubules would increase, while if case 2) were true, distance between cells would remain similar, but distance between cells from different tubules would decrease.

**Figure 6:**
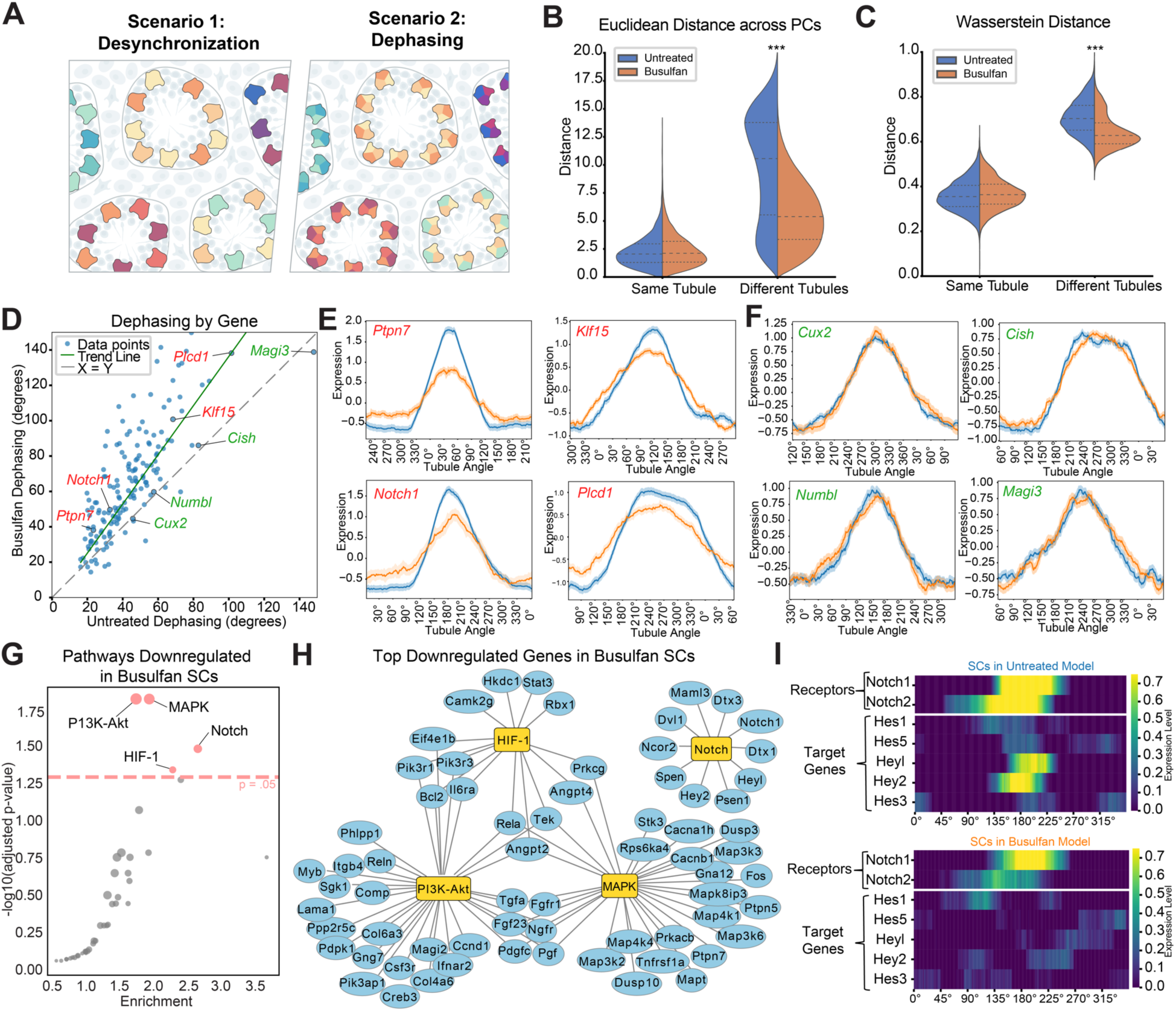
Sertoli cells independent of germ cells retain intra-tubular synchronization but inter-tubular desynchronization characterized by gene level dephasing and downregulation across major signaling pathways. **(A)** Schematic illustrating two possible scenarios of tubule level gene desynchronization after germ cell ablation. Scenario 1: Sertoli cells within the same tubule desynchronize from one another to cause a mixed expression profile for tubule. Scenario 2: Single cells within a tubule dephase and show mixed expression profiles typically expressed orthogonally. **(B-C)** Violin plots showing Euclidean (B) and Wasserstein (C) distances between Sertoli cells within the same tubule and across different tubules in untreated and busulfan-treated conditions. *** indicates p < 0.001. **(D)** Quantification of gene dephasing between busulfan and untreated conditions, measured in degrees of lag (Tubule Angle) until autocorrelation drops below 0.80 (Methods). The gray dashed line represents no dephasing, while the green trend line shows the overall shift towards increased dephasing in the busulfan treated Sertoli cells. Example dephased genes are highlighted in red, while example genes with no dephasing are highlighted in green. **(E-F)** Gene expression profiles across tubule angles for selected genes in untreated (blue) and busulfan-treated (orange) conditions. (E) shows genes with showing evidence of dephasing, while (F) shows genes with no dephasing. **(G)** Volcano plot of pathway enrichment analysis for downregulated genes in busulfan-treated Sertoli cells. Red dots indicate significantly enriched pathways in the Kegg Pathways database. **(H)** Network representation of top down-regulated genes in busulfan-treated Sertoli cells in pathways found to be significantly downregulated. **(I)** Heatmaps showing expression of Notch receptors and target genes across tubule angles in WT (top) and busulfan-treated (bottom) Sertoli cells. Untreated Sertoli cells showed activation of Notch, evidenced by expression of target genes while Busulfan-treated Sertoli cells did not. *See Figure S9 for additional data*.

Our results showed that while *intra*-tubular distance between Sertoli cells remained similar, *inter*-tubular distance between Sertoli cells decreased significantly(Fig. 6B, C). Thus, while Sertoli cells in the busulfan condition remain synchronized to a similar degree, displaying similar expression profiles with their neighbors of the same tubule, each tubule has less distinct expression profiles than in the untreated condition. In support of this, when analyzed at a single cell level, Sertoli cells from the busulfan condition occupy a space within the circular topology of untreated Sertoli cells in PC space (Fig. S8E,G).

To better understand this dephasing, we quantified the level of dephasing by computing the correlation decay with time shifts (Methods). Across 208 genes we found significant dephasing, with an average of ∼32 degrees. We also noticed a trend where genes that typically had a very narrow temporal expression profile were less dephased compared to other genes with broader expression profiles (Fig. 6D,E).

To quantify changes in gene expression at a pathway level we identified the top 200 downregulated genes and found they were significantly enriched in genes related to the PI3K-Akt, MAPK, HIF-1, and Notch signaling pathways (Fig. 6G). At a network level, while PI3K-Akt, MAPK and HIF-1 signaling pathways share many connections, the Notch pathway seems independent (Fig. 6H, Fig S9). In untreated Sertoli cells *Notch1* was expressed around Stage VII, along with respective ligands, such as *Dll1*, *Dll3*, *Jag1*, *Jag2*, being expressed at the same in many germ cell types. Notably, this co-expression is also accompanied by the expression of transcription factors tied to Notch activation such as *Hes1, Hes5, Heyl, Hey2,* and *Hes3* (Fig 6I). However, while *Notch1* was expressed in Sertoli cells from the busulfan condition, they lacked expression of *Hes1, Hes5, Heyl, Hey2,* and *Hes3* providing evidence that lack of germ cell communication resulted in decreased expression across the Notch signaling pathway (Fig 6I). In support of this, ligands were expressed around the same phase in germ cells that would activate Notch signaling pathways in Sertoli cells (Fig S10A,D,G). Thus, the decrease in distance between tubule profiles along the cyclic transcriptional profile for Sertoli cells in the busulfan condition compared to untreated is likely due to the lack of communication with germ cells, causing dephasing across gene expression profiles and downregulation of communication dependent pathways (Fig. S11J).

### Sertoli cells, independent of germ cells, display an innate retinoic acid cycle and a connected cycle of transcription factors

The discovery of a cyclic transcriptional profile in Sertoli cells – even without germ cells – suggests an innate oscillatory gene program. Since retinoic acid (RA) signaling is known to synchronize spermatogenesis and influence its timing, we analyzed the transcriptional dynamics of RA metabolic components ^43,46^. We found that Sertoli cells in the busulfan condition possess an innate RA cycle, displaying a distinct cyclic pattern in the expression of genes involved in RA metabolism (Fig. 7A,B). This intrinsic RA cycle in Sertoli cells can be divided into three main phases: an early stage (0° - 120°) characterized by balanced production (high *Aldh1a1*) and degradation (high *Cyp26b1*), suggested an overall low RA level; a middle stage (120° - 225°) marked by high RA production (high *Aldh1a2*) and no degradation, leading to high RA levels; and a late stage (225° - 360°) showing no RA production, high degradation (high *Cyp26a1* and *Cyp26c1*), and preparation for the next cycle via increased *Rdh10* expression for Retinal formation (Fig. 7A,B). Importantly, these 3 phases could be explained by biochemical feedback loops that would drive a limit cycle^43^ (Fig 7A). In the middle stage, elevated RA levels inhibit further RA production and drive degradative enzymes, leading to the late stage with low RA. This transitions into the early stage, where the rates of RA degradation and production are balanced causing a buildup of oxidized metabolites. These metabolites then inhibit further degradation and activate enzymes that produce more RA, returning the cycle to the middle stage. This Sertoli cell RA cycle persists even in the absence of germ cells, suggesting it is autonomous. However, in the intact testis, this baseline rhythm is likely modulated by the presence of differentiating germ cells that produce RA along Stage VII - VIII providing additional feedback via retinoic acid receptors on Sertoli cells ^43,47^. The cyclic nature of RA metabolism in Sertoli cells may also explain why external RA administration can synchronize the spermatogenic cycle ^48,49,43,46^. Exogenous RA likely pushes the system towards the middle stage state, where RA levels are naturally highest, effectively resetting all Sertoli cells to this phase and allowing subsequent synchronized progression through the cycle. This mechanism could also facilitate synchronization of Sertoli cells within a local tubule environment, consistent with literature ^43,50^.

**Figure 7:**
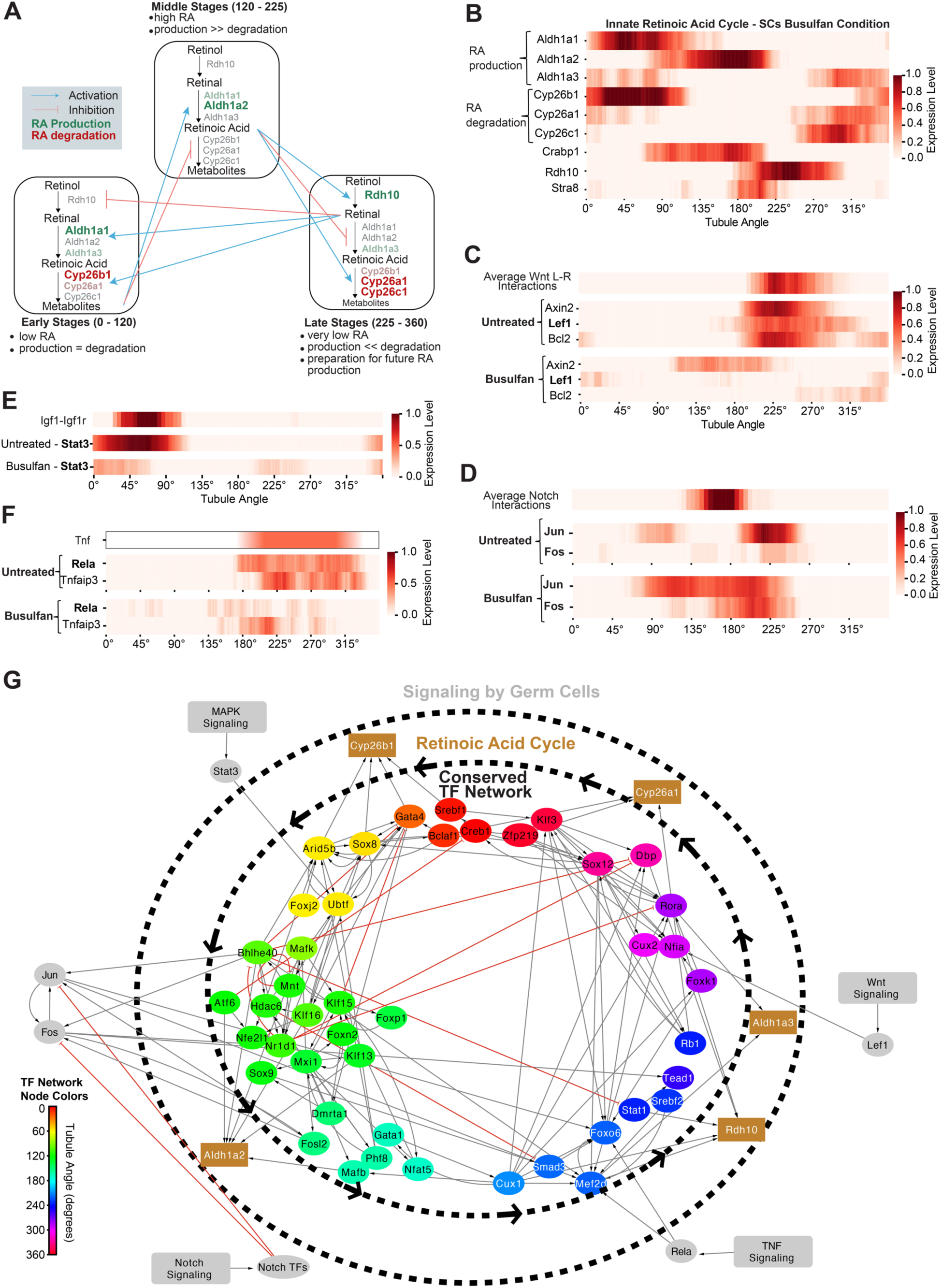
Sertoli Cells present an innate retinoic acid cycle and transcriptional machinery for oscillations that coordinates with signaling from germ cells. **(A)** Schematic representation of the Sertoli cell innate retinoic acid cycle, divided into early, middle, and late stages. **(B)** Heatmap showing the expression of genes involved in retinoic acid metabolism across Tubule Angles in busulfan-treated testes. **(C)** Heatmaps showing expression of average Wnt signaling interactions in the untreated condition, and downstream expression of canonical Wnt target transcription factors (*Axin2, Lef1, Bcl2*) in the untreated and busulfan conditions. While expression of target TFs are expressed in untreated, they are not expressed in the busulfan condition. **(D)** Heatmaps showing expression of AP1 transcription factors Jun and Fos across tubule angles in untreated and busulfan-treated testes in relation to Notch interactions present in the untreated condition. AP1 transcription factors seem to be inhibited by Notch in the untreated condition, but lacking inhibition in the busulfan condition. **(E)** Heatmap showing ligand-receptor interaction between *Igf1-Igf1r* resulting in expression in *Stat3* in the untreated condition. The busulfan-treated Sertoli cells lack similar Stat3 expression. **(F)** Heatmaps showing expression of TNF signaling-related genes across tubule angles in untreated and busulfan-treated testes. Untreated Sertoli cells show expression of downstream TNF signaling genes including *Rela, Tnfaip3,* while Sertoli cells from the busulfan condition do not. Based on literature, *Tnf* is expressed from stage VII - XI in germ cells^74^. **(G)** Conserved TF network was inferred via the Scenic pipeline^41^ using connections conserved in the busulfan condition and untreated condition (Supplementary Table 2). Connections to retinoic acid genes were inferred using busulfan condition Sertoli cells, which were screened for retinoic acid metabolism genes. Connections to signaling by germ cells were identified by identifying signaling pathways(Notch, TNF, Wnt, and MAPK) with visibly absent downstream targets in the busulfan condition Sertoli cells. Many of these absent TFs showed connections to the conserved network in the untreated condition, suggesting how germ cell signaling can coordinate with the innate TF network. *See Figure S11 for additional data*.

While retinoic acid is a key cycling component, it is likely that other networks contribute to the Sertoli cells maintaining their transcriptional oscillation. Using the SCENIC pipeline^41^, we found a sparsely connected network of transcription factors (TFs) with cyclic activation that is conserved in both untreated and busulfan-treated Sertoli cells (Fig. 7G, Supplementary Table 2). We observed features in the network including sparse connectivity, presence of feedback loops, and a hierarchical structure. The structure of this transcription factor network shares some features with known oscillatory systems, suggesting a potential mechanism that could contribute to cyclic gene expression^51–53^. The preservation of this network in both untreated and busulfan-treated conditions hints at its potential role in maintaining cyclic gene expression patterns. Supporting the functional relevance of this network we found a coherent association between transcription factor activity and target gene activity between the conditions; synchronized transcription factors predominantly targeted genes that also remained synchronized, while TFs that were not synchronized predominantly targeted genes that were not synchronized (Fig S11H-I). For example, *Klf15,* which remains synchronized across conditions, also has synchronized target genes *Foxp1*, *Mxi1*, and *Nfat5* (Fig. 7G, Supplementary Table 2).

Among the transcription factors identified in the network, we observed several known repressive interactions that could contribute to generating a limit cycle (Fig. 7G). These include repression of *Dbp, Stat1, and Gata4* by *Bhlhe40* (also *Dec1*) ^54–57^, repression of *Rora*, *Dbp, Bhlhe40* by *Nr1d1 (*also *Rev-Erbɑ*) ^58–60^, repression of *Smad3* by *Hdac6* ^61^, repression of *Gata4* by *Klf15* ^62^, and negative autoregulation of *Mnt* ^63^. Future studies could explore different parameters, such as regulatory strengths, time decays, degradation rates, and post-translational dynamics, needed to generate an oscillatory network.

Many of the genes involved in these repressive interactions (*Bhlhe40, Nr1d1, Rora, Dbp*) are known components of the circadian clock machinery. However, our observed cycle is unrelated to intra-day dynamics since all Sertoli cells are observed from a cross section spanning many tubules captured at a single time point, eliminating daily circadian or hormonal fluctuations as drivers for this network. The presence of circadian clock components in this longer-term oscillatory network suggests a repurposing of these molecular mechanisms to generate and maintain cycles on a different timescale. This repurposing is supported by the antiphase relationship between *Bhlhe40/Nr1d1* and *Dbp/Rora*, mirroring dynamics in the circadian rhythm^56,60^. Similar repurposing of circadian clock genes for non-circadian processes have been observed in other biological cycles such as the 3 week hair follicle cycle in mouse dorsal skin^64^ and the 12-hour rhythms in the liver ^65,66^.

To understand how this conserved transcription factor(TF) network could interact with the identified retinoic acid(RA) cycle, we analyzed the TFs for potential regulation of RA cycle genes. Several TFs showed enriched binding motifs in the RA cycle gene promoters and matched the expression phase (Fig. 7G). We also identified two TFs known to be inducible by retinoic acid: *Bhlhe40* (also known as Sensitive to retinoic acid 13, *Stra13*) and *Mafb* ^67–70^. These genes peaked between 150° and 200°, coinciding with the expected peak of retinoic acid levels (Fig. 7A,B). These temporal alignments suggest a feedback mechanism between the TF network and the RA cycle, potentially contributing to the robustness and coherence of Sertoli cell rhythms, while retinoic acid remains the major regulator of tubular level synchronization^43^.

Finally, we investigated whether signaling pathways, including those we identified as downregulated (Fig 6G,H), influence the TF network through germ cell communication. We found germ cell communication would reinforce the network through interactions involving the Wnt, Notch, TNF, and MAPK signaling pathways. In untreated control, interactions across these signaling pathways lead to expression of known canonical TFs that interact with the network in a phase consistent manner. However, in the Busulfan condition, these TFs are not expressed or lack coherent expression. These TFs included *Lef1*(Wnt pathway, Fig 7C), *Hes/Hey* TFs(Notch Pathway, Fig 6I), *Rela* (TNF pathway, Fig 7F), and *Stat3* (MAPK pathway, Fig 7E) respectively. Notably, AP-1 transcription factors *Jun and Fos* showed distinct expression patterns in relation to Notch signaling (Fig 7D). In untreated conditions, Jun expression occurs before and after Notch activation, while Fos activity follows it. However, in the busulfan condition, both Jun and Fos are expressed throughout typical Notch-active phases (Fig. 7D), aligning with models where Notch signaling indirectly represses AP-1 transcription factor activity ^71,72^. These observations indicate that germ cell-derived signals can enhance the robustness and synchronization of the transcriptional oscillation in Sertoli cells by reinforcing the TF network through signaling pathways. Moreover, these communication pathways may provide a mechanism for germ cells to modulate the timing of spermatogenesis, which could further explain observations where donor germ cells alter the length of the seminiferous epithelial cycle ^73^.

## Discussion

In this study, we present a comprehensive spatial and temporal map of spermatogenesis at a single-cell resolution, revealing novel insights into the organization and regulation of this complex process. Our findings not only provide a detailed characterization of the seminiferous epithelial cycle but also uncover a previously unrecognized role for Sertoli cells as innate oscillators coordinating with spermatogenesis.

Our application of spatial transcriptomics via RNA SeqFISH+ leverages the unique spatial geometries of seminiferous tubules, where each tubule cross-section represents a distinct time point, to infer temporal dynamics. This approach overcomes limitations of previous studies that relied on dissociated cells, enabling us to reconstruct the temporal dynamics of spermatogenesis with unprecedented resolution. The circular transcriptional profile we observed in the gene expression space of tubule cross-sections precisely recapitulates the known stages of the seminiferous epithelial cycle, validating our approach and providing a quantitative framework for mapping temporal dynamics using spatial geometries. This method of bridging spatial geometries to temporal dynamics has broad application in various biological systems where time is encoded in space. For instance, in other cyclical biological processes, such as hair follicle cycle, bone remodeling, and intestinal epithelial turnover, where different stages are spatially segregated, our approach could help reconstruct temporal dynamics from spatial snapshots ^75–79^. Moreover, this approach can be generalized to infer pseudotimes for non-cyclical processes in tissues with spatially organized developmental or functional gradients.

A striking finding of our study is the observation that Sertoli cells exhibit a cyclic transcriptional profile closely synchronized with germ cell development. The fact that Sertoli cells were the only somatic cell type to show such synchronization further emphasizes their unique position in the regulation of spermatogenesis. This is consistent with early studies that have identified anatomical changes in Sertoli cells over the seminiferous epithelial cycle ^28,29^. However, it is important to note why previous sequencing studies did not fully capture this phenomenon; those using scRNA-seq and microarray, provided insights into stage-dependent expression^1,26,27^ but were limited in their ability to directly correlate transcriptional dynamics with germ cell development dynamics due to the dissociation process or limited resolution. Furthermore, Sertoli cells are significantly underrepresented in scRNAseq datasets due to the difficulties with the dissociation process^1,4^. Our spatial transcriptomics approach, in contrast, preserves this crucial spatial context allowing us to observe how Sertoli cell transcriptional profiles change in synchrony with germ cell development.

Importantly, we found that this cyclic transcriptional profile is maintained autonomously in Sertoli cells, even in the absence of germ cells, suggesting an intrinsic oscillatory program that may coordinate with the synchronized development of germ cells, reminiscent of other biological oscillators such as the segmentation clock in somitogenesis or the circadian rhythm^80^. However, while the cyclic transcriptional profile remained, there was significant downregulation of signaling pathways such as PI3K-Akt, MAPK, and Notch. Thus, the absence of germ cells suggests that while a core oscillator is autonomous, its fine-tuning and robustness depend on reciprocal signaling with germ cells. This interplay between cell-autonomous oscillations and cell-cell communication is reminiscent of the coupling between cellular clocks that maintains coherent rhythms at the tissue and organism level ^81–83^.

Our findings finally reveal two cyclic processes in Sertoli cells that may underlie the mechanism for the observed cyclic transcriptional profile. First, the intrinsic RA cycle, characterized by three distinct phases, serves to synchronize Sertoli cells within a tubule even in the absence of germ cells. This cycle may also explain the synchronizing effect of exogenous RA on spermatogenesis observed in previous studies ^10,43,46,50^. Second, we identified a sparsely connected network of transcription factors displaying cyclic activation. Importantly, both cycles persist in Sertoli cells even in the absence of germ cells, suggesting they form the basis of an autonomous oscillatory system. However, the presence of germ cells likely provides additional modulation, refinement, and reinforcement of these processes, resulting in the precise orchestration of spermatogenesis observed in vivo. This is supported by the downregulation of signaling pathways in Sertoli cells in the absence of germ cells, and transcription factors only expressed in the presence of germ cells that would target genes in the oscillatory network, such as *Lef1, Rela, Hes/Hey,* and *Stat3.* This would also explain why transplantation experiments where rat germ cells supported by mouse Sertoli cells still developed at the rate of the rat seminiferous epithelial cycle^73^. While somatic cells in various stem cell niches are known to influence stem cell behavior ^84–86^, the discovery of an autonomous, cyclic transcriptional program in Sertoli cells represents provides a new perspective on how somatic niche cells might coordinate with the complex timing of stem cell behavior, and provides insights into the general principles of biological oscillators and tissue organization.

### Limitations of the Study

While our study captured the cycle of the Sertoli cell, several important questions remain. First, without the presence of differentiating germ cells we cannot capture if the period of oscillation differs from the normal seminiferous epithelial cycle. Second, our RNA-based measurements do not capture potential dynamic regulation through post-translational modifications. Third, while our transcription factor network is inferred from binding motifs and single-cell expression data, direct experimental validation of these connections is still needed. Further studies should investigate how the Sertoli cell cyclic transcriptional profile interacts with germ cells in transplantation experiments where the period of the seminiferous epithelial cycle changes ^27,73,87^. While Sertoli cells may present as baseline rhythm, this rhythm may even be overridden by specific cues such as retinoic acid and other signaling from germ cells ^43,46^. These insights could have far-reaching implications for reproductive biology, especially as sequencing studies suggest that many genes associated with idiopathic human non-obstructive azoospermia, accounting for ∼15% of male infertility cases, were predominantly expressed in Sertoli cells ^88,89^.

## Acknowledgements

We thank I. Strazhnik for help with the figures; N. McMillan for helping with the intraperitoneal mouse injections; K.L. Colon for the preprocessing pipeline^90^; H. Amhrein for showcasing our data in his interactive explorer; J.Fox for help with mice handling. We also thank all Cai Lab members for the insightful comments and discussions.

Work in LC’s laboratory was supported by the Allen Discovery Center for Cell Lineage Tracing (to L.C), training grants including the NIH T32GM008042 (to UCLA-Caltech MSTP), and the David Geffen Medical Scholarship (to A.C).

## Author Contributions

Conceptualization: A.C., L.C., B.D.S., S.Y. Experiments and Data Collection: A.C. Data Analysis: A.C. Interpretation of Results: A.C., L.C., S.Y., B.D.S. Writing: A.C., S.Y., B.D.S., L.C. Supervision: L.C.

## Declaration of interests

L.C. is a cofounder of Spatial Genomics, Inc. The remaining authors declare no competing interests.

## Data and Code availability

Scripts used for preprocessing seqFISH+ images can be found at https://github.com/CaiGroup/pyfish_tools. Source and processed data will be available on Figshare after completion of the review process. Processed data can be interactively explored via our web interface https://woldlab.caltech.edu/ci2-celltiles/MouseSpermatogenesis_AC/ (best in Firefox).

## Supplementary Tables

**Supplementary Table 1:** A list of genes included in this study, including barcoding information for each gene.

**Supplementary Table 2:** A summary transcription factors and their conserved targets between the untreated and busulfan condition.

## Methods

### Gene Selection

An initial panel of mouse transcription factors and signaling pathway genes (total 3073 genes) was selected from the Kegg Pathways Database. This initial panel included genes from major signaling pathways (e.g. MapK, Notch, Wnt, Jak-Stat etc.) and all transcription factors in mouse. Expression levels for these genes were then analyzed using bulk-RNA seq expression data from the Mouse Encode transcriptome data covering 30 mouse tissues (GEO: GSE36025)^91^. As a generalized mouse tissue pool, genes with FPKM levels >1000 across many tissues were removed from the panel. In addition, genes for known housekeeping genes (e.g. *Eef2*) were removed. The final barcoded panel includes 2622 genes covering >1300 transcription factors and >1300 signaling genes. A full list of genes is provided in Supplemental Table 1. For these barcoded genes the average FPKM per hybridization was estimated to be ∼1300 in mouse testis. Based on previous experiments, this was an ideal dot density to avoid optical crowding.

To supplement our gene pool for studying spermatogenesis we included an additional pool of marker genes that would be visualized using smFISH for stem cells, spermatogonia, spermatocytes, spermatids, Sertoli and Leydig cells. In total, the final experiments covered 2638 genes, including 2622 barcoded genes and 16 genes using smFISH. For the busulfan condition, we further added additional genes for visualization using smFISH related to retinoic acid metabolism. The full list of genes is provided in Supplementary Table 1.

### RNA seqFISH+ encoding strategy

To spatially resolve mRNA profiles for 2622 genes in the mouse testis we used a modified version of RNA seqFISH+ encoding scheme. The 2622 were split into two pools, one containing signaling genes (1322 genes) while the other contained all transcription factors (1300 genes). Each pool was encoded using 8-pseudocolor bases with 4 rounds of barcoding including one-error correction round, which can accommodate up to 512 genes, in one fluorescent channel (635 nm), and 1536 genes across 3 fluorescent channels (640nm, 561nm, 488nm) (Supplementary Table 1). An additional 16 genes for mRNA marker genes were encoded as a non-barcoded seqFISH scheme. In total 2638 genes were measured in the untreated wildtype condition.

In the busulfan condition, the same two pools containing signaling pools and transcription factors was used, but a different group of non-barcoded genes was included to include retinoic acid metabolism genes, for a total of 2653 genes (Supplementary Table 1).

### Primary-probe design

To obtain probe sets for >2600 different genes, 35-nucleotide (nt) sequences of each gene were extracted, using the exons from within the coding region. The masked genome and annotation from the University of California Santa Cruz (UCSC) were used to look up the gene sequences. Probe sequences were required to have GC content within the range 45–65%. Any probe sequences that contained five or more consecutive bases of the same kind were dropped. Any genes that did not achieve a minimum number of 24 probes were dropped. A local BLAST query was run on each probe against the mouse transcriptome to ensure specificity. BLAST hits on any sequences other than the target gene with a 15-nt match were considered off targets. ENCODE RNA-seq data across different mouse samples were used to generate an off-target copy-number table. Any probe that hit an expected total off-target copy number exceeding 10,000 FPKM was dropped to remove housekeeping genes, ribosomal genes and very highly expressed genes. To minimize cross-hybridization between probe sets, a local BLAST database was constructed from the probe sequences, and probes with hits of 17 nt or longer were removed by dropping the matched probe from the larger probe set.

Once 35nt sequences for each probe was obtained, we assigned barcodes to each gene randomly across the potential codebook. Because of the large number of genes, random assignment created roughly equal expression in each hybridization round. These barcodes corresponded to pseudocolors that would be read out using specific readout probes. The corresponding reverse complements for the readout probes, effectively representing a specific barcode, were appended as overhangs to either side of the primary probe. While each gene received a barcode of 4 pseudocolors (described in RNA seqFISH+ encoding strategy), each probe received a total of 6 readout sites corresponding to the 4 pseudocolor barcode. Thus, for any individual probe 2 of the pseudocolors had two readout sites, while the other 2 pseudocolors had one readout site. By varying which readout sites were repeated across all the probes there were 1.5 readout sites per pseudocolor per probe on average for any given gene.

### Readout-probe design and synthesis

Readout probes of 15-nucleotide length were designed as previously introduced^17^. In brief, the probe sequences were randomly generated with combinations of A, T, G or C nucleotides, with a GC content in the range of 40–60%. To validate the specificity of the generated readout sequences, we performed a BLAST search against the mouse transcriptome. To minimize cross-hybridization of the readout probes, all probes with 10 contiguously matching sequences between the readout probes were removed. The reverse complements of these readout probe sequences were included in the primary probes according to the designed barcodes. The fluorescently-labeled readout probes (Integrated DNA Technologies) that can bind to the readout sequences on the primary probes were conjugated in-house to Alexa Fluor 647–NHS ester (Invitrogen A20006), Cy3B–NHS ester (GE Healthcare PA63101), or Alexa Fluor 488– NHS ester (Invitrogen A20000) as described before^17,18^ or directly purchased (Integrated DNA Technologies).

### Primary-probe construction

Primary probes were ordered as oligoarray complex pools from Twist Bioscience and were constructed as previously described with some modifications^17^. In brief, limited PCR cycles were used to amplify the designated probe sequences from the oligo complex pool. Then, the amplified PCR products were purified using the QIAquick PCR Purification Kit (28104; Qiagen) according to the manufacturer’s instructions. The PCR products were used as the template for in vitro transcription (E2040S; NEB) followed by reverse transcription (EP7051; Thermo Fisher). After reverse transcription, the probes were subjected exonuclease I(M0293S; NEB) to digest any unused remnant primers, while the complete RNA-DNA hybrids would remain intact. Then, the single-stranded DNA (ssDNA) probes were alkaline hydrolyzed with 1 M NaOH at 65 °C for 15 min to degrade the RNA templates, followed by 1 M acetic acid neutralization. Finally, to clean up the probes, we performed ethanol precipitation to remove stray nucleotides, phenol– chloroform extraction to remove protein, and used SPRIselect beads (B23317, Beckman) to remove any residual nucleotides and phenol contaminants. The probes were stored at −20 °C until use.

### SeqFISH+ Imaging

SeqFISH+ imaging was carried as described previously^17,90^. In brief, the flow cell of the sample was connected to the automated fluidics system. Then the region of interest (ROI) was registered using nuclei signals stained with 7.5μg ml^−1^ DAPI (D8417; Sigma). Each serial hybridization buffer contained three unique sequences with different concentrations of 15-nt readouts conjugated to either Alexa Fluor 647 (50 nM), Cy3B (50 nM) or Alexa Fluor 488 (100 nM) in EC buffer (10% ethylene carbonate (E26258; Sigma), 10% dextran sulfate (D4911; Sigma), 4× SSC and 1:100 dilution of SUPERase In RNase inhibitor(AM2694; Invitrogen). The 100 μl of serial hybridization buffers for 70 rounds of seqFISH+ imaging (including smFISH genes) with a repeat for round 1 (in total 71 rounds) was pipetted into a 96-well plate. During each serial hybridization, the automated sampler moves to the well of the designated hybridization buffer and moves the 100 μl hybridization buffer through a multichannel fluidic valve (EZ1213-820-4; IDEX Health & Science) to the flow cell (requires ∼25 μl) using a syringe pump (63133-01, Hamilton Company). The serial hybridization solution was incubated for 20min at room temperature. After serial hybridization, the sample was washed with ∼300 μl of 10% formamide wash buffer (10% formamide and 0.1% Triton X-100 in 2× SSC) to remove excess readout probes and non-specific binding. Then, the sample was rinsed with ∼300 μl of 4× SSC supplemented with a 1:1,000 dilution of SUPERase In RNase Inhibitor, before being stained with DAPI solution (7.5 μg ml^−1^ of DAPI, 4× SSC, and a 1:1,000 dilution of SUPERase In RNase inhibitor) for ∼15 s. Next, an anti-bleaching buffer solution made of 10% (w/v) glucose, 1:100 diluted catalase (Sigma C3155), 0.5 mg ml^−1^ glucose oxidase (Sigma G2133), 0.02 U μl^−1^ SUPERase In RNase inhibitor and 50 mM pH 8 Tris-HCl in 4× SSC was flowed through the samples. Imaging was done with a microscope (Leica DMi8) equipped with a confocal scanner unit (Yokogawa CSU-W1), a sCMOS camera (Andor Zyla 4.2 Plus), a 63× oil objective lens (Leica 1.40 NA) and a motorized stage (ASI MS2000). Lasers from CNI and filter sets from Semrock were used. Snapshots across a single z slice per FOV across 647-nm, 561-nm, 488-nm and 405-nm fluorescent channels. After imaging, stripping buffer (55% formamide and 0.1% Triton-X 100 in 2× SSC) was flowed through for 1 min, followed by an incubation time of 3 min before rinsing with 4× SSC solution. In general, the 15-nt readouts were stripped off within seconds, and a 3-min wash ensured the removal of any residual signal. The serial hybridization, imaging and signal extinguishing steps were repeated for 64 rounds. Then, staining buffer for segmentation After all RNA SeqFISH+ imaging rounds, concanavalin-A (conA) conjugated to Alexa Fluor 647 (ThermoFisher C21421) was flown into the tissue using the SeqFISH fluidics system at 50ug/mL in 1X PBS for 5 minutes. Afterwards, the sample was washed 3 times with 55% Wash Buffer for 5 minutes each time to wash out residual conA. The tissue was then imaged again but with a shorter 300ms exposure time in the 647 channel. The integration of automated fluidics delivery system and imaging was controlled by a custom written script in μManager.

### Image Processing and Decoding of SeqFISH data

Image analysis was performed as previously described^90^ with some modifications.

#### Image alignment

Image alignment was performed either using phase correlation on DAPI-stained images for each field of view, or using scale-invariant feature transform. The segmentation hybridization round was used as a reference for estimating translational shifts in the x and y direction.

#### Image preprocessing

A 5×5 high-pass Gaussian filter removed residual background, followed by a 3×3 low-pass filter to mitigate hot pixels and enhance spot Gaussian profiles for improved 2D fitting. Image intensities were normalized across channels and serial hybridizations using 80-99.999% percentile clipping and 0-1 rescaling.

#### Dot detection

Sub-pixel spot centroids were identified using DAOStarFinder from Astropy, performing fast 2D Gaussian fits. FWHM was optimized for optimal spot calling. Features recorded include flux, peak amplitude, sharpness, symmetry (bilateral to four-fold), and Gaussian fit symmetry. Additional features, like total spot area, were obtained using a 7×7 bounding box and local adaptive thresholding with a Gaussian kernel.

#### Decoding

Super-resolved, mapped spots undergo SVM-embedded, feature-based symmetrical nearest neighbor decoding. An SVM model with RBF or polynomial kernel filters false spots based on characteristics. Spots receive likelihood scores for true barcode correspondence. A radial search (0.75 - 1.5 pixels) across barcoding rounds scores spots on distance, intensity, and size. Best spots are chosen for each round, assigning overall codeword and ambiguity scores. Barcodes undergo parity checks, spot set consistency filtering (≥3 appearances), and overlap resolution. Unused spots are resubmitted for up to 2 additional decoding rounds. Decoded spots are sorted by codeword score and subsampled to calculate FPR. Spots with FPR ≤5% are used to generate the final gene-by-cell matrix for downstream analysis.

#### SVM training

Quick-pass decoding labels true and fake spots. The classifier activates with 500-500,000 fake spots. True spots are down-sampled to match fake spot count. Data is split 80% for training, 20% for validation. Features are normalized using MinMax Scaler. GridSearchCV with 8-fold cross-validation tunes C, gamma, and degree parameters for polynomial or radial-basis function kernels. Test set performance is evaluated using training data scaling parameters.

#### False Positive rate

False Positive Rate, or FPR was determined as following:

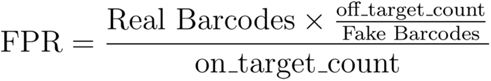

Where:

● Real Barcodes: Number of gene-coding barcodes in the codebook
● Fake Barcodes: Number of non-coding (empty) barcodes
● off_target_count: Number of decoded fake barcodes
● on_target_count: Number of decoded real barcodes

Code Availability: Scripts used for pre-processing seqFISH images can be found at https://github.com/CaiGroup/pyfish_tools.

### Cell Segmentation

After all RNA SeqFISH+ imaging rounds, concanavalin-A (conA) conjugated to Alexa Fluor 647 (ThermoFisher C21421) was flown into the tissue using the seqFISH fluidics system at 50ug/mL in 1X PBS for 5 minutes. Afterwards, the sample was washed 3 times with 55% Wash Buffer for 5 minutes each time to wash out residual conA. The tissue was then imaged again but with a shorter 300ms exposure time in the 647 channel.

To computationally identify cell masks, a Cellpose 2.0 model was trained using >1000 cells for each experiment^20,21^. The models were trained to identify cell masks using both the nuclear DAPI stains as well as the conA membrane stains. Using these trained models, we used Cellpose to obtain our final cell masks which were contracted by 2-4 pixels to further increase specificity. Finally, decoded RNA transcripts were assigned to individual cells based on whether they fell within the 2-dimensional cell masks.

### Tubule Segmentation

Since the tubules were readily distinguishable in the stitched images of testis cross-sections and were limited in number, they were manually segmented using FIJI’s ROI Manager. Each tubule was demarcated as a distinct ROI and assigned a unique integer identifier. After all the tubules were added to the ROI Manager for a given testis cross-section, a labeled mask image was generated and exported for each tubule cross section. In this image, each pixel value corresponds either to the integer identifier of a specific tubule or to zero if no tubule was present at that location. These labeled mask images were subsequently imported into Python as numpy arrays. These arrays were then used as overlay masks to identify and isolate the corresponding spatial regions in the AnnData object for tubule-level analysis or tubule identification.

### Coverslip functionalization

Coverslips were cleaned with a plasma cleaner on a high setting (PDC-001, Harrick Plasma) for 5 min, followed by rinsing with 100% ethanol three times, and heat-dried in an oven at >90 °C for 30 min. Next, the coverslips were treated with 100 μg μl^−1^ of poly-D-lysine (P6407; Sigma) in water for >5 h at room temperature, followed by three rinses with water. The coverslips were then air-dried and kept at 4 °C for no longer than a few days.

### Mice

All animal care and experiments were carried out in accordance with Caltech Institutional Animal Care and Use Committee (IACUC) and NIH guidelines. Adult 5-8 week old wild-type mice C57BL/6J were used for the experiments.

### Testis tissue extraction and processing

In brief, mice were euthanized via asphyxiation using CO2 followed by cervical dislocation and decapitation. The mouse testis were removed from the mouse within 5 minutes of death and immediately flash frozen in OCT medium using an isopentane bath that was cooled with surrounded liquid nitrogen. These flash frozen testis were stored at −80 °C until ready to be sectioned. Each experiment was conducted using a different mouse.

10 micrometer sections were cut using a cryotome and immediately placed on functionalized coverslips. Then thin tissue slices were then fixed using 4% Paraformaldehyde(28908; ThermoFisher) in 1X PBS for 10 minutes at room temperature. After fixation the slides were gently washed with 1X PBS and placed in 1X PBS baths for 10 minutes to remove residual PFA and further quenched by 100mM Tris-HCl, pH 7.4 for 10 minutes. Slides were finally rinsed again with 1X PBS. Slides were then permeabilized by placing into 70% ethanol overnight at - 4°C. Afterwards, the tissue was further permeabilized using 8% SDS(AM9822; Invitrogen) in 1X PBS for 15 minutes. 1X PBS was used to wash away any residual SDS followed by washed with 70% ethanol. The samples were then placed in 4X SSC (15557036, Thermo Fisher) overnight.

Samples were hybridized with primary probes in the hybridization buffer (35% formamide, 10% dextran sulfate and 2X SSC) at 37 °C with a concentration of ∼1.4nM/probe for 3 days. After hybridization the samples were washed with 55% WB (55% formamide and 0.1% Triton-X 100 in 2× SSC) and incubated for another 30 minutes at 37 °C to reduce any non-specific binding. The sample was washed with 4X SSC several times and placed into the microscope holder for imaging.

### Busulfan Treatment

10mg of busulfan (ThermoFisher J61348) was dissolved in 1mL DMSO and subsequently added to 9mL of DPBS to make a 1mg/mL stock solution. This stock was sterilized with a 0.2uM filter and subsequently injected intraperitoneally in 5 week old C57BL/6 mice. To completely ablate differentiating germ cells a concentration of 30mg/kg or 45mg/kg was injected. Both doses caused complete ablation of differentiating germ cells, consistent with dosing response literature^45^. The mice were then housed in sterile housing due to the potential immunosuppressive properties and busulfan. After 30 days from injection, the testes were collected from the mice euthanized via asphyxiation using CO2 followed by cervical dislocation and decapitation.

### Modified Moran’s I

Each cross section of the testis consists of seminiferous tubule cross sections organized spatially. To assess whether there is global spatial organization in the stages of these tubules, we employed a spatial autocorrelation analysis. While Moran’s I is a standard tool for such analyses, it has limitations when applied to cyclic datasets. Namely, Euclidean distances fail to accurately measure adjacency between points, e.g. 360° = 0°. To address these limitations, we developed a modified version for cyclic parameters, *I_cyclic_*, similarly developed for neural imaging data^92^.

One way of defining distance for circular data is to treat each angle/phase as a point on the unit circle of the complex plane. Thus, distance between two angles would be computed using a 2-argument arctangent function with a full range from -180 to 180:

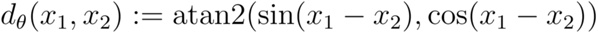

To compute the average angle, *x̄* can be calculated by computing the average component of sin, *S̄* and the average component of cosine, *C̄*.

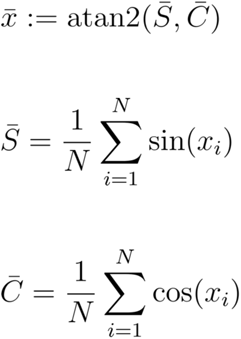

Thus, the full equation for the modified Moran’s I for cyclic parameter, *I*_cyclic_, is:

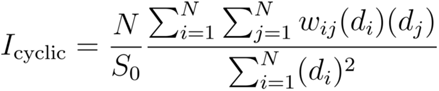

Where:

*N*: the number of tubules

*d_i_* and *d_j_* : the distance between angle values *x_i_* and *x_j_*, respectively, and the mean angle *x̄*

*S*_0_ : sum of all spatial weights 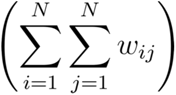

### Gene expression preprocessing and clustering

The python package Scanpy (version 1.10)^93^ was used to process the cell by gene matrices. For the single cell analysis, counts were normalized to 1000, log normalized, and scaled using the pp.normalize_total(), pp.log1p(), and pp.scale() function respectively. Batch correction was performed prior to scaling using the built-in combat module in scanpy^94^. The functions pp.pca(), pp.neighbors() and tl.umap() was used for dimensionality reduction and cluster identification resulting in 30 clusters. The distribution of clusters across samples and imaged positions was examined and 4 clusters showed a strong position bias caused by technical issues during imaging rounds resulting in decreased barcodes decoded that included specific hybridization rounds. These positions were removed altogether from one dataset. Differentially expressed genes between clusters were identified using tl.rank_genes_groups() with the standard wilcox statistical testing. The same process was used for the tubule level analysis, except the counts were normalized to 10,000.

### Determining order of tubules and Tubule Angle

The order of tubules in both untreated and busulfan conditions was determined using the de-novo cyclic pipeline in scPrisma^23^. Prior to running the pipeline, counts were normalized and log-scaled. The cyclic order was reconstructed using the ‘reconstruction_cyclic’ function and the function was run for 500-1000 iterations to ensure convergence and stability of the cyclic ordering. To ensure robustness of the cyclic ordering, we repeated the pipeline multiple times for each dataset and compared the resulting cyclic orders for consistency.

After establishing the cyclic order, we converted these orders to Tubule Angles evenly distributed across 0 - 360 degrees. This conversion from order to Tubule Angle is justified and appropriate based on our demonstration of random organization of tubules across testis cross-sections, which we established using spatial autocorrelation analysis (see Figure S4 and Modified Moran’s I section). The absence of global spatial organization in tubule stages across the testis cross-section enables us to confidently represent the cyclic order as a continuous variable mapped onto a circular space.

### Mapping to Tubule Stages

The frequency and time scale of the stages of the seminiferous epithelial cycle have been well characterized by sampling across 3,000 random tubule cross sections from 12 wildtype mice^11^. Our sample size of 337 random tubule cross sections in untreated wildtype testis was sufficiently large to enable accurate mapping of the order/Tubule Angle across these previously characterized frequencies. We validated the stages of tubules primarily through analysis of cellular composition (Figure 2, Figure S6), complemented by examination of DAPI morphology. The stages we determined aligned precisely with the expected cellular composition and DAPI morphology for each stage of the seminiferous epithelial cycle. This mapping process allowed us to confidently associate each Tubule Angle with a specific stage of the seminiferous epithelial cycle, providing a robust framework for analyzing gene expression patterns across the progression of spermatogenesis.

### Population Dynamics

To analyze the distribution of different cell types across the seminiferous epithelial cycle, cell type information was extracted from the AnnData object, creating separate dataframes for each unique cell type. We developed a custom function to bin the data into 6-degree intervals across the 360-degree cycle and calculate circular moving averages using a window size of 5 bins. This approach accounted for the circular nature of the data, ensuring continuity at the 0/360-degree boundary. A custom plotting function was then used to visualize the distribution of cell types, where the x-axis represents the cycle angle (0-360 degrees) and the y-axis shows the average count of cells per tubule with 95% confidence intervals displayed as shaded areas.

### SCENIC analysis

The python implementation of the SCENIC pipeline^41^ was used for regulatory network reconstruction providing insight into *cis-*regulatory insight between transcription factors and target gene expression. First, GRNBoost2, a gradient boosting algorithm, was used to infer gene regulatory networks from the expression data at the single cell level. This step was performed using the SCENIC CLI tools. Afterwards cisTarget was used to predict regulons. This step involves identifying enriched transcription factor binding motifs in the regulatory regions of the target genes. We used three different motif databases for this analysis specific for mouse and covering different genomic regions (500bp upstream, 10kb centered on TSS, and 5kb centered on TSS). For motif annotations, we used the file ’motifs-v9-nr.mgi-m0.001-o0.0.tbl’, which provides information about mouse transcription factor motifs used in the analysis. The ’mgi’ in the filename indicates that it uses Mouse Genome Informatics (MGI) nomenclature, ensuring mouse-specific annotations. The results from this analysis were used to identify key transcription factors and infer target genes.

The core transcription factor network conserved between untreated and the busulfan condition was identified by retaining conserved connections between TFs that were inferred in both the untreated and the busulfan condition.

### Quantifying synchronization

To compare the synchronization between somatic cells and the seminiferous epithelial cycle at a topological level, we employed a computational approach using the scPrisma algorithm^23^. We first created separate AnnData objects for germ cells and the somatic cell type of interest (e.g., endothelial cells), aggregating gene expression data at the tubule level. For somatic cells that were outside tubule masks, they were assigned the tubule that was closest. We then used scPrisma’s cyclic reconstruction function to independently determine the ordering of tubules based on gene expression patterns in both germ cells and somatic cells. This process was repeated 31 times using parallel processing to ensure robustness, with each iteration performing the reconstruction for both germ cells and somatic cells. The resulting orders from each run were saved separately for both cell types. The circular correlations between the orderings were then used to assess the degree of synchronization between the somatic cells and the seminiferous epithelial cycle (Figure 4A,B). To quantify significance, we used a bayesian statistics approach using PyMC3 to compare the mean difference in correlations between each somatic cell type and the null models. The likelihood for each group was modeled as a normal distribution and the probability that the difference falls within a region of practical equivalence (ROPE) was evaluated and reported as our p value.

To quantify gene expression at the gene level for each somatic cell type, we developed a computational pipeline that analyzed expression patterns in relation to the spermatogenic cycle. For each gene, we calculated the moving average of expression across the cycle using a window size of 5 and bin size of 6 degrees. To assess the significance of these expression patterns, we performed 1000 permutations for each gene, randomly shuffling the cell angles to generate a null distribution. We then calculated 99% confidence intervals from these permutations to define the expected noise level. Real expression patterns were compared against these noise intervals to determine if a gene exhibited significant cyclic expression (Figure 4F). This process was parallelized using a job array such that each gene was assigned a separate computational node. The analysis was performed separately for each somatic cell type (e.g., peritubular cells, Sertoli cells). Genes were considered to have significant cyclic expression if their real expression pattern exceeded the noise confidence intervals for more than 50 contiguous degrees of the cycle.

To quantify gene dephasing between untreated and busulfan-treated samples, we developed a computational pipeline that analyzed expression patterns across the spermatogenic cycle. First, we preprocessed the data using Scanpy, normalizing to 1000 counts per cell, applying log-transformation, and performing batch correction via combat. We then ordered genes based on their dynamic expression patterns and extracted expression data for both untreated and busulfan-treated samples. To smooth the data, we applied a circular moving average with a window size of 50 degrees. We then employed Fourier transform-based autocorrelation to assess the dephasing of gene expression patterns. The degree of dephasing for each gene was estimated by determining the lag at which the autocorrelation dropped below a threshold of 0.80. This process was performed for both untreated and busulfan-treated samples. The lag at which the autocorrelation dropped to 0.80 was plotted for both busulfan and untreated genes (Figure 6D). Thus, a gene with a larger lag to reach the threshold autocorrelation in the busulfan condition compared to untreated would be found to be dephased. This approach allowed us to quantify the extent to which Busulfan treatment disrupted the cyclic expression patterns of individual genes.

### Gene and Pathway enrichment analysis

Gene Ontology analysis for gene expressed dynamically in Sertoli cells along the seminiferous epithelial cycle was performed using the *GOATOOLS* Python library. Mouse gene symbols were converted to Entrez Gene IDs using the *mygene* package. The GO database (go-basic.obo, dated 2023-07-27) and mouse gene association files were downloaded from the Gene Ontology Consortium. Enrichment analysis was run using the GOEnrichmentStudyNS() function. The background gene set included only genes in our study, and p values were adjusted using the Benjamini-hochberg procedure. Only GO terms that were statistically significant, having an adjusted p value < .05, were reported.

To identify pathways and genes altered in busulfan-treated samples compared to untreated, we employed a multi-step computational approach. The data was preprocessed including normalizing counts to 1000 counts per cell, applying log-transformation, and performing batch correction. We then calculated moving averages of gene expression across the spermatogenic cycle using a 50-degree window. To quantify changes in gene expression patterns, we computed the Kolmogorov-Smirnov statistic between untreated and busulfan-treated samples for each gene. We focused on the top 200 genes with the highest KS statistic for further analysis. Using the clusterProfiler R package, we performed pathway enrichment analysis on these genes, utilizing a custom-filtered set of KEGG pathways focused on signaling cascades. We used a conservative background set that included only genes in the KEGG pathways and also present in our experiment. After enrichment analysis we employed the Benjamini-Hochberg method for multiple testing correction. Pathways with an adjusted p-value < 0.05 were considered significantly enriched. This approach allowed us to identify both individual genes and broader pathways disrupted by the lack of germ-cell communication.

### Ligand-Receptor Analysis

Ligand-receptor interaction analysis was performed using gene expression data from germ cells and Sertoli cells. A comprehensive list of ligand-receptor pairs was compiled from CellTalkDB^95^. The interaction scores were computed as the product of the normalized ligand and receptor expression values when expression of ligands in germ cells and receptors in Sertoli cells occurred within the same tubule. Thus, interaction scores were calculated across all tubules and these were then standardized using z-score normalization. To identify dynamic interaction patterns, the standardized scores were smoothed using a moving average with a window size of 5 bins, each representing 6 degrees. These custom functions are available in the FigShare folder. This analysis allowed for the identification of spatially-regulated ligand-receptor interactions between germ cells and Sertoli cells throughout the seminiferous epithelial cycle.

---

**Figure S1:**
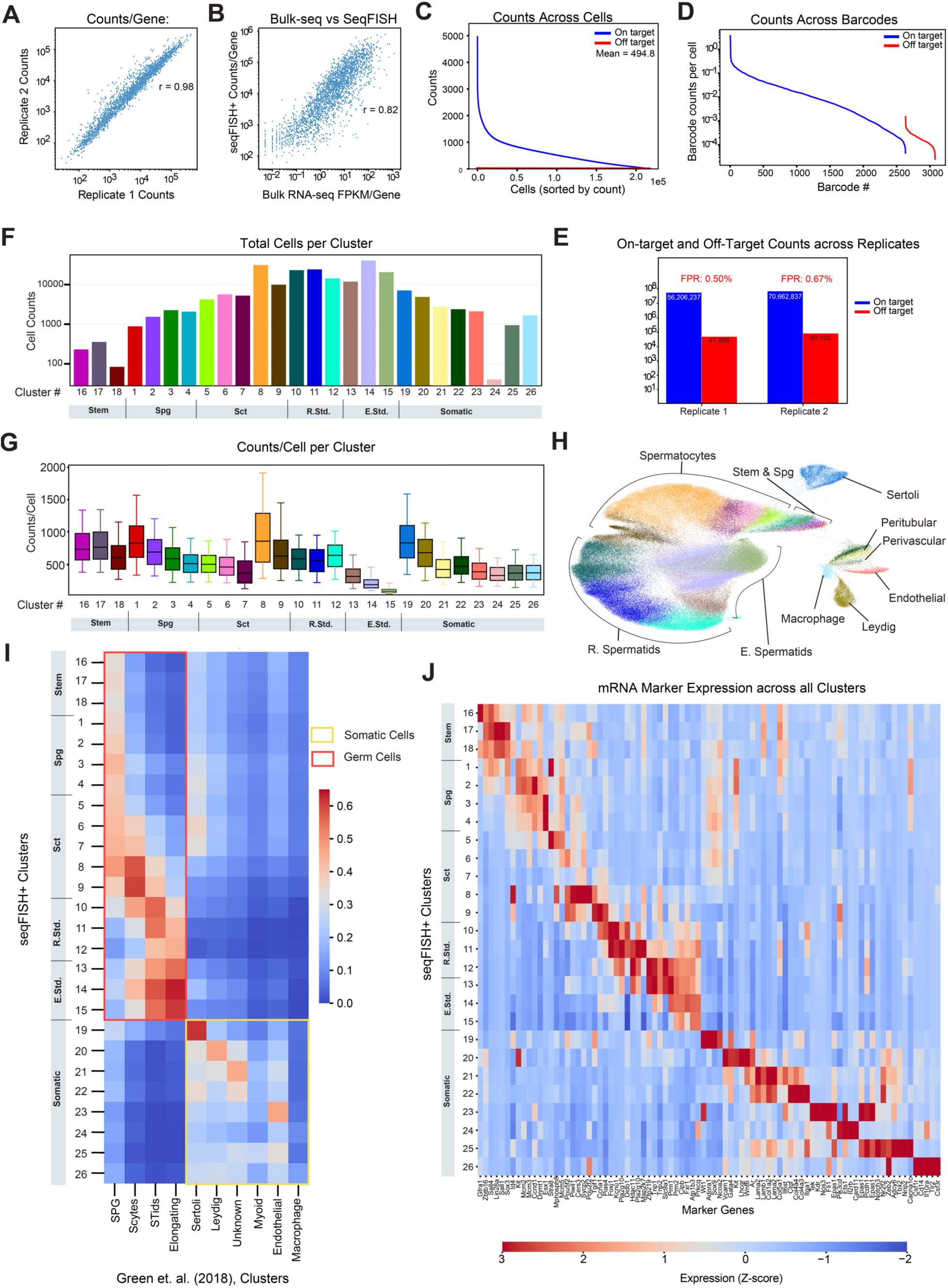
Validation and characterization of SeqFISH+ measurements in adult mouse testes. **(A)** seqFISH+ counts per gene were highly reproducible, with a Pearson correlation of 0.98 across the two replicates. **(B)** seqFISH+ counts per gene were consistent with measurements from Bulk RNA-seq with a Pearson correlation of 0.82. **(C)** Visual representation of on- and off-target barcode counts in each cell. Off-target barcodes were those barcodes that did not map to any gene in the probe pool. Of the 3072 possible barcodes, 2622 mapped to genes while 450 mapped to unused barcodes and were considered off-target barcode counts. The median false positive rate for each cell was ∼0.00% while the mean false positive rate was ∼0.60%. **(D)** Counts across all barcodes. On target barcodes showed significantly higher barcode counts/cell than off-target barcodes. **(E)** Histogram showing on-target barcodes vs off-target barcodes across both wildtype/untreated replicates. The false-positive rate, calculated by normalizing the available off-target barcodes in the codebook, was 0.50% in replicate 1 and 0.67% replicate 2. In replicate 1, the total number of on-target and off-target counts were 56,206,237 and 47,868 respectively. In replicate 2, the total number of on-target and off-target counts were 70,662,837 and 80,102, respectively. **(F,G)** Total cells per cluster and counts/cell per cluster are shown. Clusters 16, 17, 18 represent Renewal Primed Stem Cells, Differentiation Primed Stem Cells, and Other Stem Cells, respectively. Clusters were ordered from 1 – 26, consistent with the order of cell types presented in Figure 1C. Specifically, Clusters 1, 2, 3, 4 represent A1-A3 Spg, A3-A4 Spg, In Spg, and B Spg, respectively. Clusters 5, 6, 7, 8, 9 represent Preleptotene, Leptotene, Zygotene, Pachytene, and Diplotene/Secondary Spermatocytes, respectively. Clusters 10, 11, 12 represent Round1-3, Round4-6, Round 7-8, respectively. Clusters 13,14,15, represent Elongating 9-10, Elongating 11-13, Elongating 14-16, respectively. Finally, Clusters 19,20,21,22, 23, 24, 25, 26 represent Sertoli, Peritubular, Peritubular 2, Leydig, Endothelial, T-Cell, Perivascular, and Macrophages, respectively. **(H)** Uniform Manifold Approximation and Projection (UMAP) of integrated seqFISH+ datasets. Cluster colors are consistent with definitions from subfigure F and G. Germ cell clusters are grouped along a continuum consistent with the path of differentiation while Somatic cells cluster separately. **(I)** seqFISH+ clusters were compared with clusters from those obtained in a single cell RNA-seq study1. Pearson correlations between cluster averages showed consistent cell-type definitions. A previously unknown cluster from the scRNA-seq dataset shows mapping to the two peritubular types. **(J)** A heatmap of the scaled expression of cell-type marker genes across all seqFISH+ clusters.

**Figure S2:**
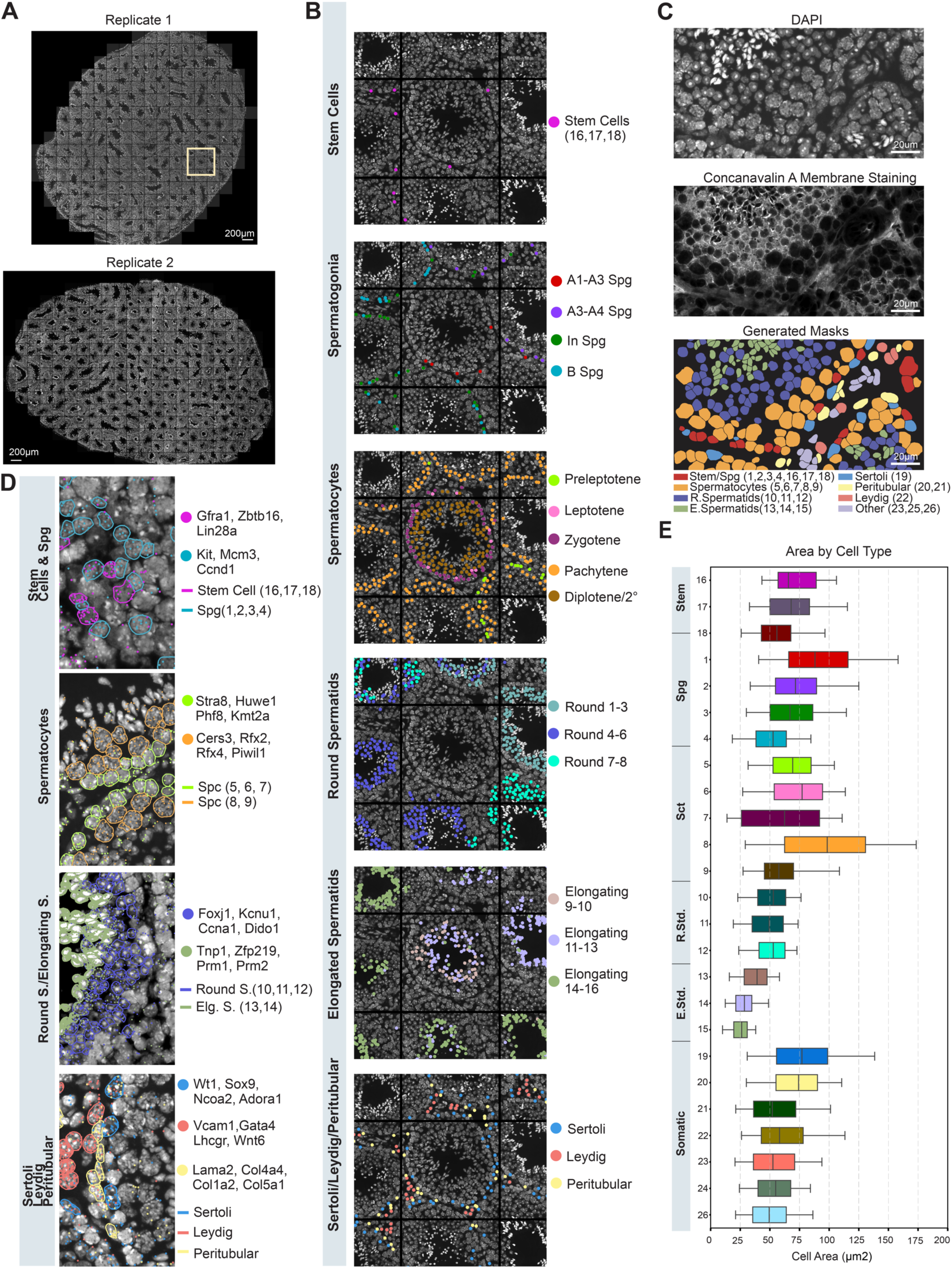
Validation of cell-types identified and spatial locations. **(A)** Stitched DAPI images of two biological replicates showing full wild-type testis cross-sections used for seqFISH+ analysis. Replicate 1 covered 271 positions while Replicate 2 covered 223 positions. Scale bar: 200 μm. **(B)** Spatial maps of identified cell types across different stages of spermatogenesis and supporting cells. From top to bottom: stem cells, spermatogonia, spermatocytes, round spermatids, elongating spermatids, and somatic cells (Sertoli, Leydig, and Peritubular cells). Each colored dot represents a cell, with the color corresponding to the cell type or stage as indicated in the legend. Spatial maps corresponded precisely with identified cell types. **(C)** Validation of cell segmentation. Top: DAPI staining showing nuclei. Middle: Concanavalin A membrane staining. Bottom: Generated masks from the using a Cellpose 2.0 custom-trained model, with colors corresponding to different cell types as indicated in the legend. **(D)** Validation of marker genes for different cell types and stages. From top to bottom: stem cells and spermatogonia, spermatocytes, round and elongating spermatids, and somatic cells. Marker genes are listed for each cell type/stage and correspond to the colored dots in the images. **(E)** Quantification of cell area by cell type, showing the distribution of cell sizes across different stages of spermatogenesis and supporting cell types. The x-axis shows cell area in μm², and the y-axis lists cell types/stages.

**Figure S3:**
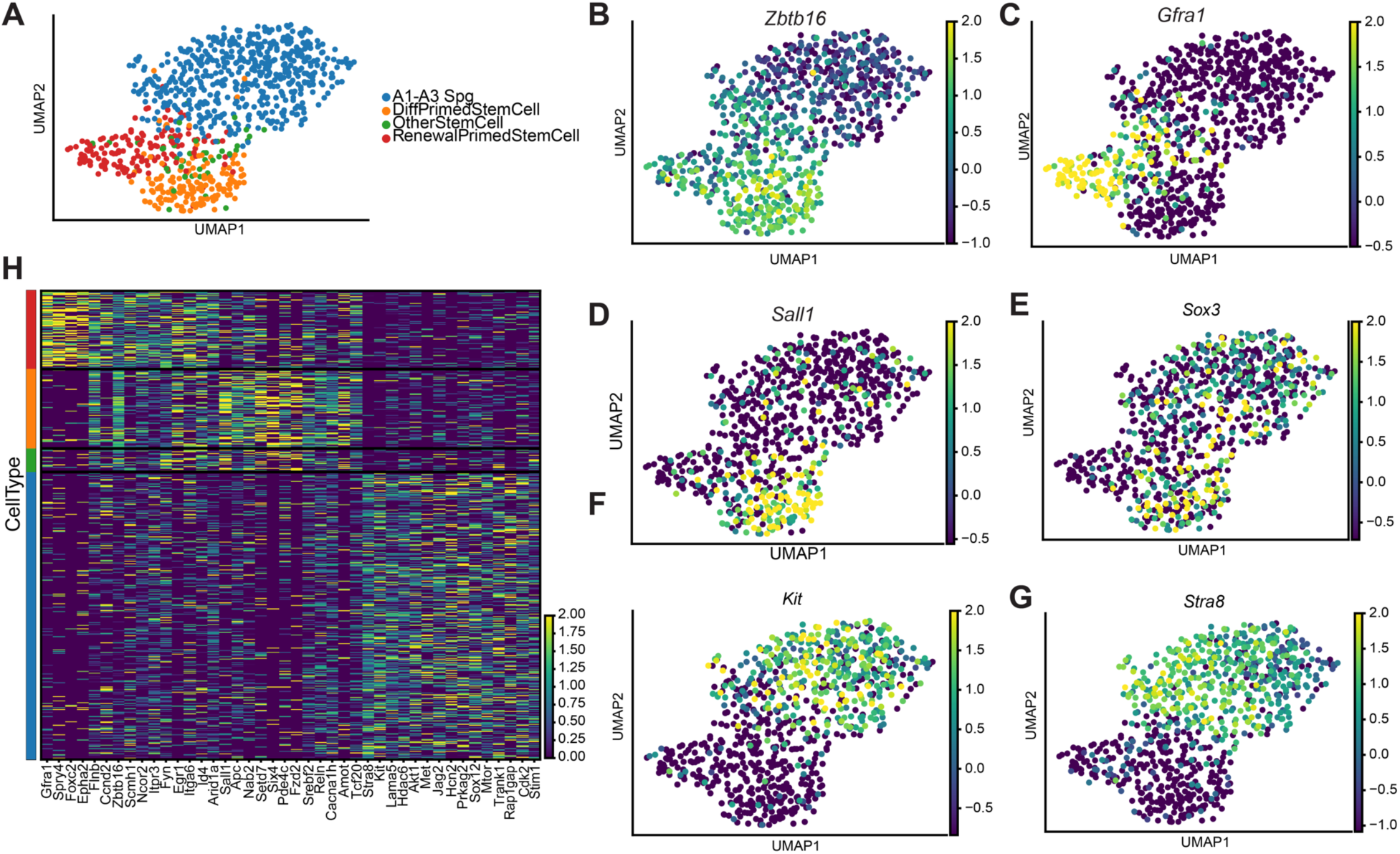
Expression profiles of Stem Cells and Early Spermatogonia. **(A)** UMAP projection of different stem cell types and early spermatogonia. **(B - G)** Marker gene expression shown in UMAP plots. *Zbtb16* is a general stem cell marker. *Gfra1* is a marker for stem cells primed for renewal. *Sall1* and *Sox3* are markers for stem cells primed for differentiation. *Sox3* shows continued expression into A1 -A3 Spg stages. *Kit* and *Stra8* mark the commitment towards the spermatogenesis differentiation pathway, expressed in A1 -A3 Spg from Stage VII - VIII. **(H)** Heatmap of additional marker genes shown across all stem cells and A1-A3 Spg.

**Figure S4:**
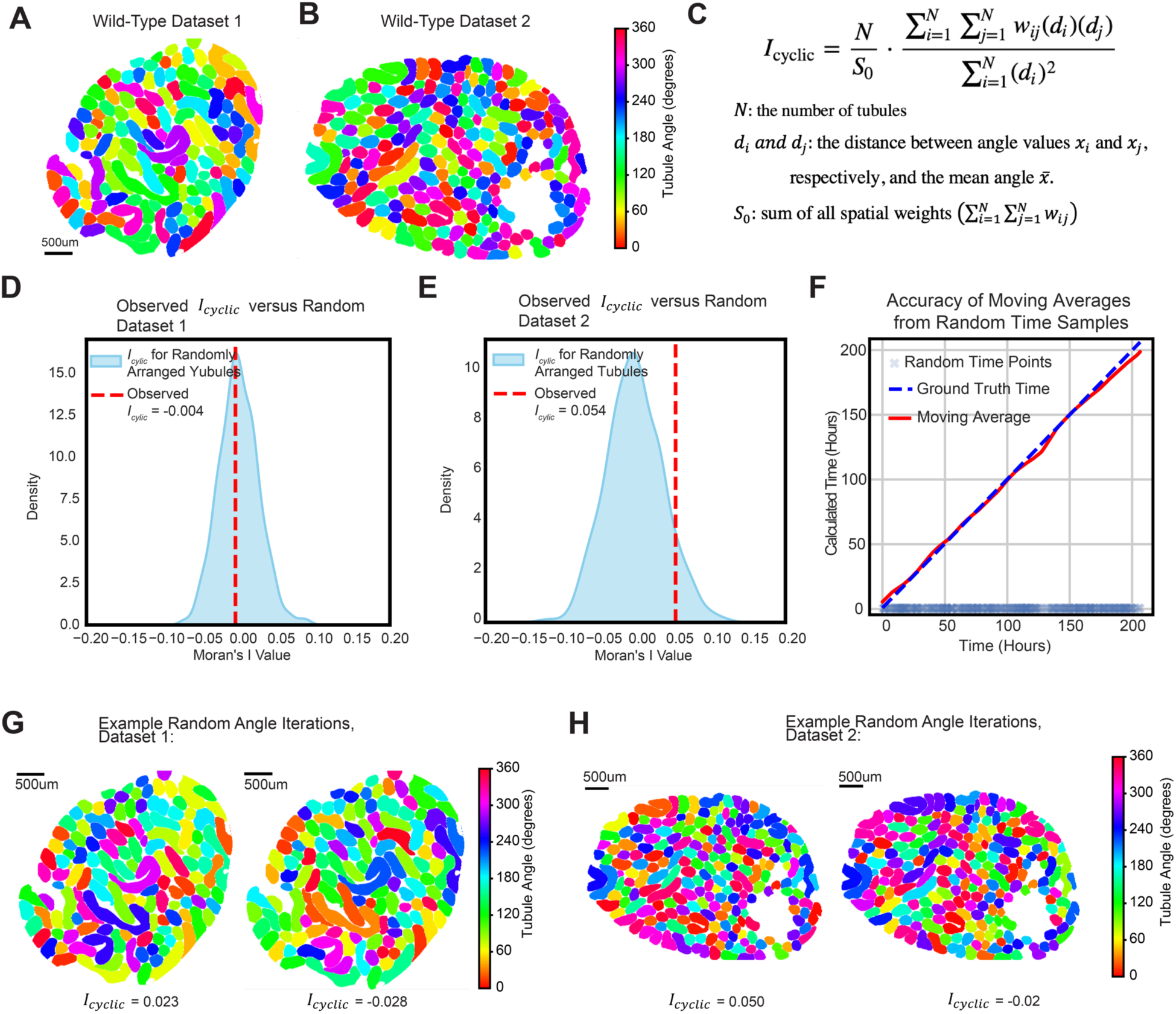
A Modified Moran’s I for Cyclic Parameters shows that Tubule Stages/Angles are organized randomly in testis cross sections. (A, **B)** Two wild-type replicates of testes are shown with tubules colored by Tubule Angle. **(C)** A modified Moran’s I, *I*_cyclic_, was developed to analyze the spatial autocorrelation for the cyclic parameter, tubule angle. Details on the derivation are provided in Methods. **(D, E)** The observed Moran’s I for each replicate was compared with the distribution of Moran’s I values if tubule angles were assigned randomly across all tubules. The observed Moran’s I fell within the distributions for random assignment. Furthermore, the values for Moran’s I were close to zero (-0.004 and 0.054, p values: 0.875, 0.167 respectively), providing strong evidence that Tubule Angles/Stages are organized randomly in testis cross sections. **(F)** Because tubule angles observed are spatially randomly organized in testes cross sections, each tubule cross section represents a random distinct time point in the seminiferous epithelial cycle. Using a windowed moving average of 30 time points, 337 random time points along the ∼208-hour seminiferous epithelial cycle can precisely recapitulate the true temporal dynamics. **(G-H)** Example cross sections with tubule angles randomized with computed *I*_cyclic_ values.

**Figure S5:**
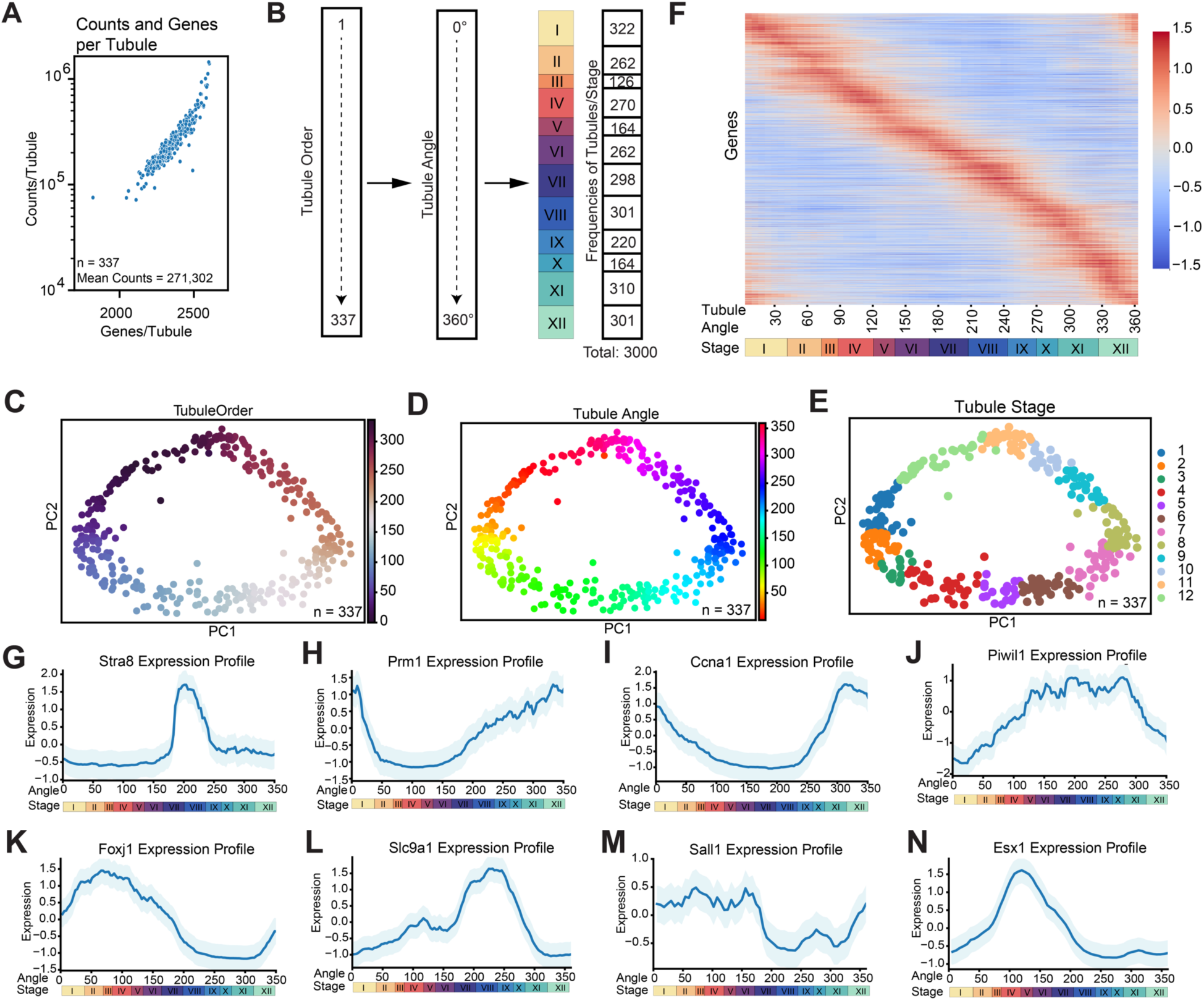
Order, Angle, and Stages for Tubules. **(A)** The counts and genes/tubule across 337 wildtype tubules are shown. On average, counts equaled 271,302 while mean genes detected was 2343. **(B)** The order of tubules was determined using the de-novo cyclic pipeline in scPrisma^23^. After establishing the cyclic order, we converted these orders to Tubule Angles evenly distributed across 0 - 360 degrees. This conversion from Order to Angle is justified and appropriate based on our demonstration of random organization of tubules across testis cross-sections, which we established using spatial autocorrelation analysis (Figure S4 and Modified Moran’s I section). Tubules Stages were then assigned based on previously characterized frequencies^11^ and validated through analysis of cellular composition and DAPI morphology (Figure 2, Figure S5). **(C-E)** Principal component plots showing Tubule Order, Tubule Angle, and Tubule Stage, respectively. **(F)** Gene expression along Tubule Angle and Stage across all Tubules. Genes were sorted by the phase of expression. **(F-M)** Show the expression patterns of *Stra8, Prm1, Ccna1, Piwil1, Foxj1, Slc9a1, Sall1,* and *Esx1* respectively. The x-axis represents the angle within the tubule (0-360°), corresponding to stages I-XII of the seminiferous epithelium cycle as indicated by the color bar below each graph. The y-axis shows the normalized gene expression level. Blue lines represent mean expression, with light blue shading indicating the 95% confidence interval. **(F)** *Stra8* is specific to retinoic acid signaling and is expressed between Stages VII - VIII, consistent with its role in initiating meiosis. **(G)** Prm1 increases in later stages (IX - XII) corresponding to high expression in elongating spermatids and consistent with its function in chromatin condensation. **(H)** Ccna1 peaks around X - XII, showing high expression in late pachytene and diplotene/secondary spermatocytes. **(I)** Piwil1 is highly expressed in mid-late pachytene spermatocytes, thus showing a peak expression between IV -XI. **(J)** Foxj1 shows high expression from XII - II, aligning with the emergence of round spermatids. **(K)** Slc9a1 is highly expressed in late round spermatids and early elongating spermatids, coinciding with expression in stages VII - X. **(L)** Sall1 is expressed in Differentiation Primed Spermatogonia, thus shows expression from I -VII, with a steep decrease thereafter. **(M)** Esx1 is highly expressed in Spermatogonia, A1 - B, aligning with expression between I - VI.

**Figure S6:**
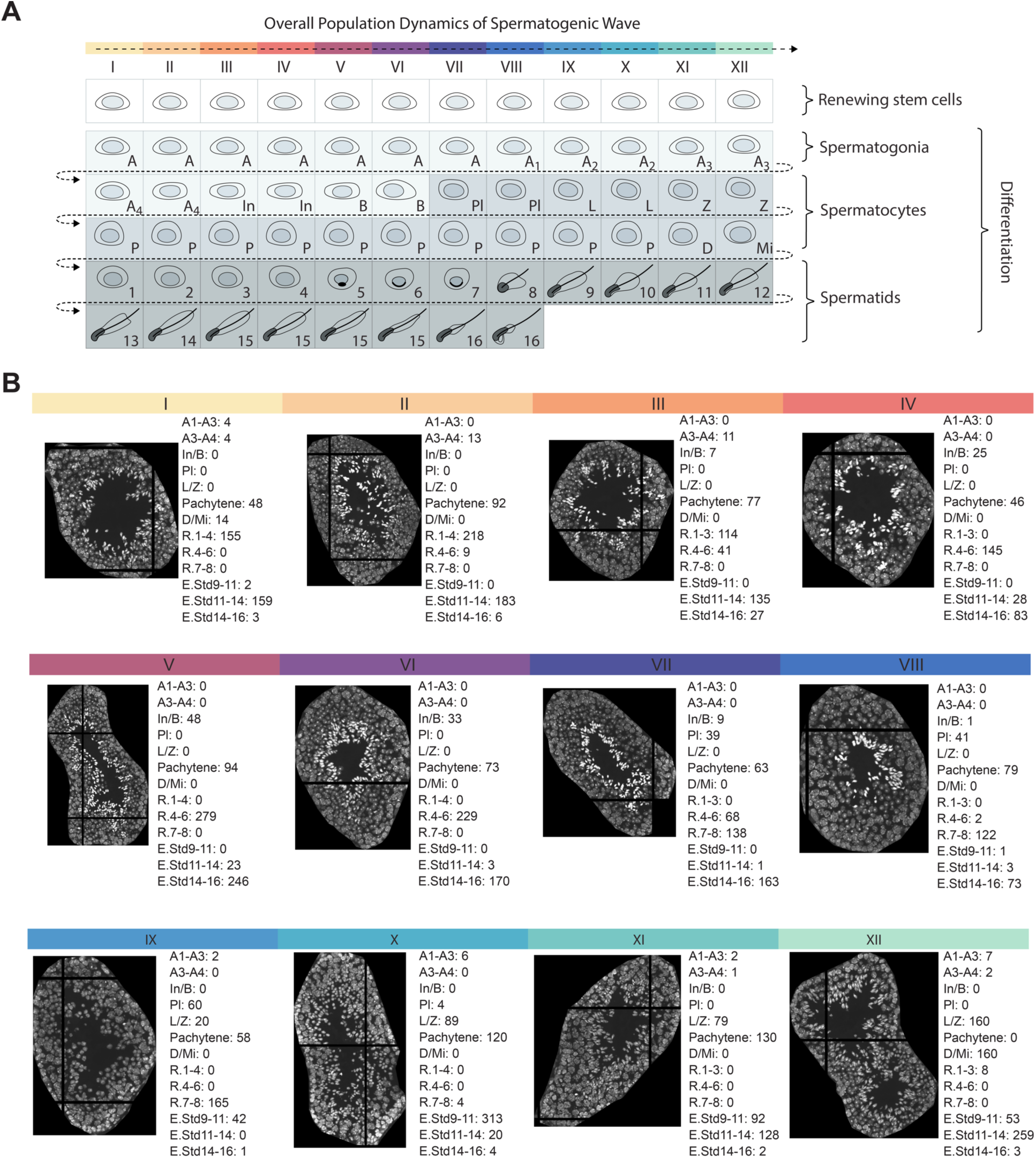
Overall Population Dynamics and Visualization of Tubule Stages and Compositions. **(A)** Overall population dynamics across stages of spermatogenic wave, each stage containing a unique composition of cell types. Abbreviations are as follows: A: Type A Spermatogonia, In: Intermediate Spermatogonia, B: Type B Spermatogonia, Pl: Preleptotene Spermatocytes, L: Leptotene Spermatocytes, Z: Zygotene Spermatocytes, P: Pachytene Spermatocytes, D: Diplotene Spermatocytes, Mi: Secondary Spermatocytes. Numerical abbreviations 1 -16 refer to rounds spermatids(1-8) or elongating spermatids(9-16). **B)** Example DAPI images of tubules identified of each stage from Replicate 1 and corresponding numbers of cell-types.

**Figure S7:**
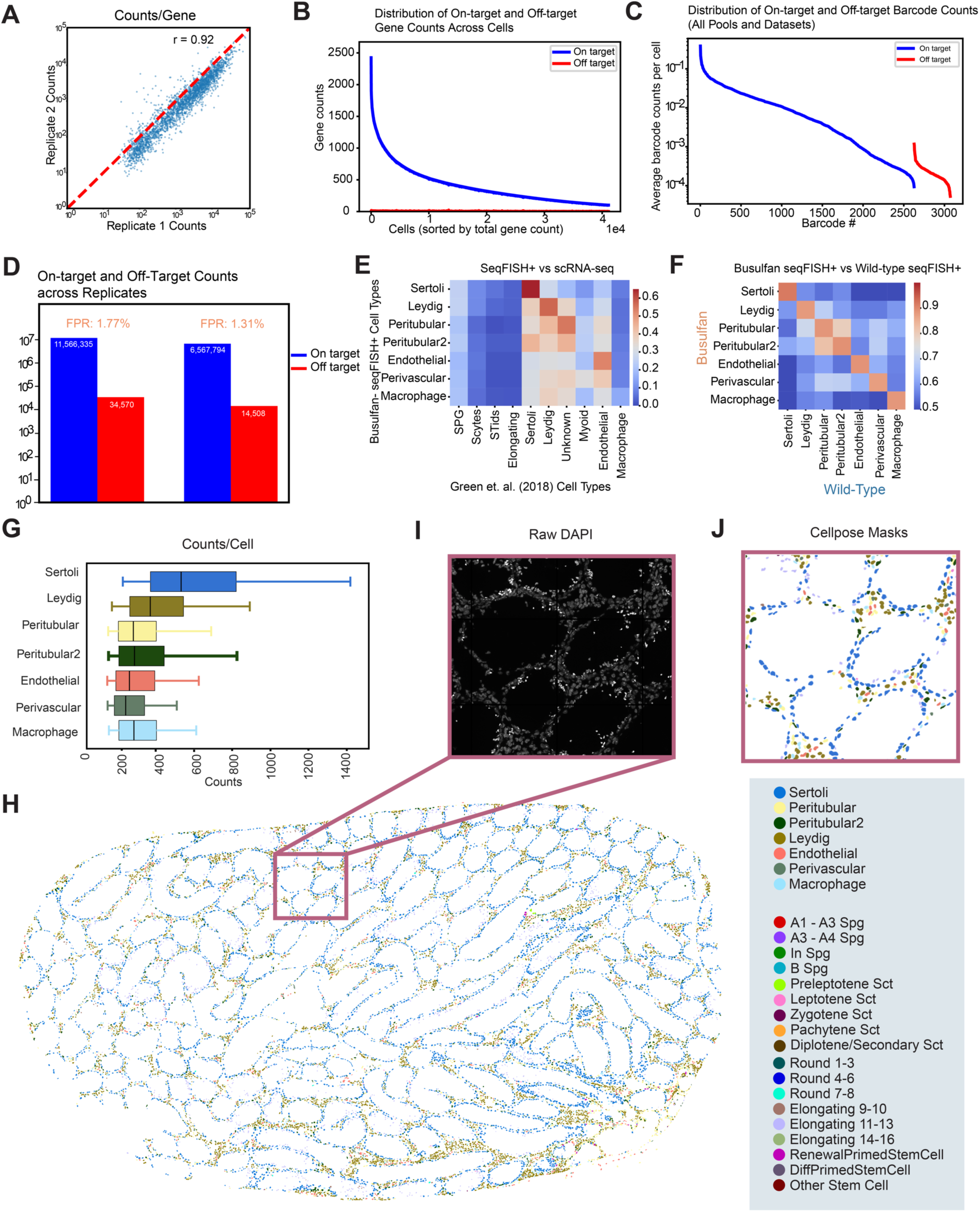
Validation and characterization of seqFISH+ measurements in adult mouse testes treated with Busulfan. **(A)** seqFISH+ counts per gene were highly reproducible, with a Pearson correlation of 0.92 across the two replicates. **(B)** Visual representation of on- and off-target barcode counts in each cell. Off-target barcodes were those barcodes that did not map to any gene in the probe pool. Of the 3072 possible barcodes, 2622 mapped to genes while 450 mapped to unused barcodes and were considered off-target barcode counts. The median false positive rate for each cell was ∼0.00% while the mean false positive rate was ∼2.60%. **(C)** Counts across all barcodes. On target barcodes showed significantly higher barcode counts/cell than off-target barcodes. **(D)** Histogram showing on-target barcodes vs off-target barcodes across both busulfan replicates. The false-positive rate, calculated by normalizing the available off-target barcodes in the codebook, was 1.77% in replicate 1 and 1.31% replicate 2. In replicate 1, the total number of on-target and off-target counts were 11,566,335 and 34,570 respectively. In replicate 2, the total number of on-target and off-target counts were 6,567,794 and 14,508, respectively. **(E)** seqFISH+ clusters were compared with clusters from those obtained in a single cell RNA-seq study^1^. Pearson correlations between cluster averages showed consistent cell-type definitions. **(F)** seqFISH+ clusters were compared between untreated and busulfan conditions showing consistent cell-type definitions across the two conditions. **(G)** Counts/cell type are shown. Whiskers represent 5 and 95 percentile. **(H)** Spatial map of cells (counts >50) represented.

**Figure S8:**
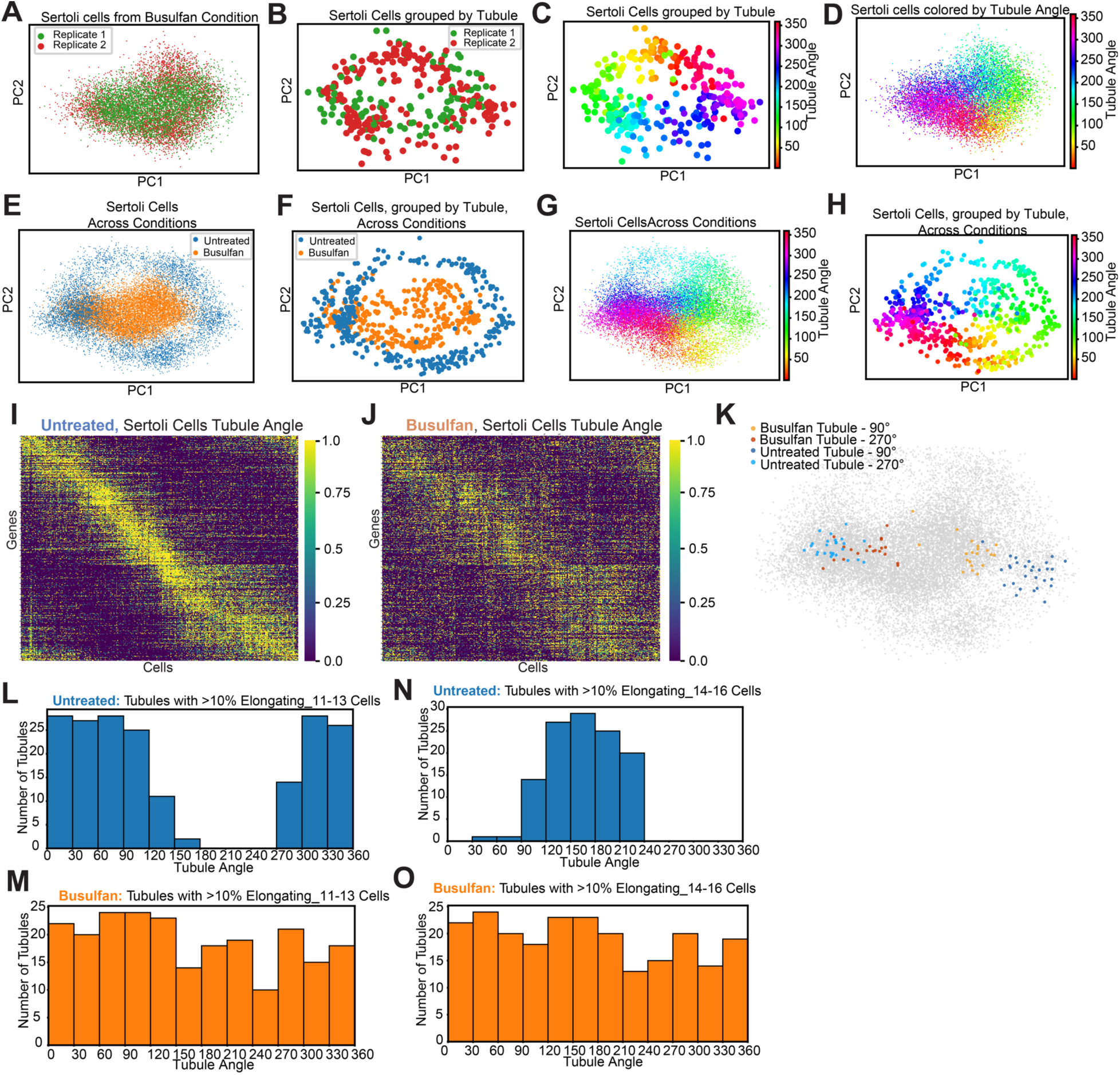
Topological analysis of Sertoli cells in Busulfan datasets. (A-D) PC level plots of busulfan condition datasets for Sertoli cells(A,D) and pseudobulked by tubule (B,C). Subfigure A,B show the datasets by replicate while subfigure C,D show the datasets colored by Tubule Angle, calculated at the tubule level independently for the busulfan condition datasets. **(E-H)** PC level plots of integrated datasets covering both busulfan and untreated conditions for Sertoli cells(E,G) and grouped by tubule (F,H). Subfigure E,F show the datasets by condition for the Single cell and grouped by tubule level respectively. Subfigure G,H show the datasets colored by Tubule Angle for the single cell and tubule level, respectively. **(I-J)** Expression data for Sertoli cells across Tubule Angle for untreated and busulfan condition, respectively. A dephasing of gene expression profiles was visible in the busulfan condition compared to the untreated condition. **(K)** The tubules at ∼90 degrees and ∼270 degrees were selected from the busulfan and untreated conditions and the cells from the constituent cells are highlighted. Across all tubules, the distance between cells in the same tubules across PC space was similar in the untreated and busulfan condition(Figure 6B, C). **(L - M)** Comparison of Tubule Angles represented by tubules with >10% Elongating Spermatids 11-13 in both untreated and busulfan condition. Since all Tubule Angles were represented in the busulfan condition, the Sertoli cells must be cycling independently of the present spermatids. **(N - O)** Comparison of Tubule Angles represented by tubules with >10% Elongating Spermatids 14-16 in both untreated and busulfan condition. Since all Tubule Angles were represented in the busulfan condition, the Sertoli cells must be cycling independently of the remnant spermatids.

**Figure S9:**
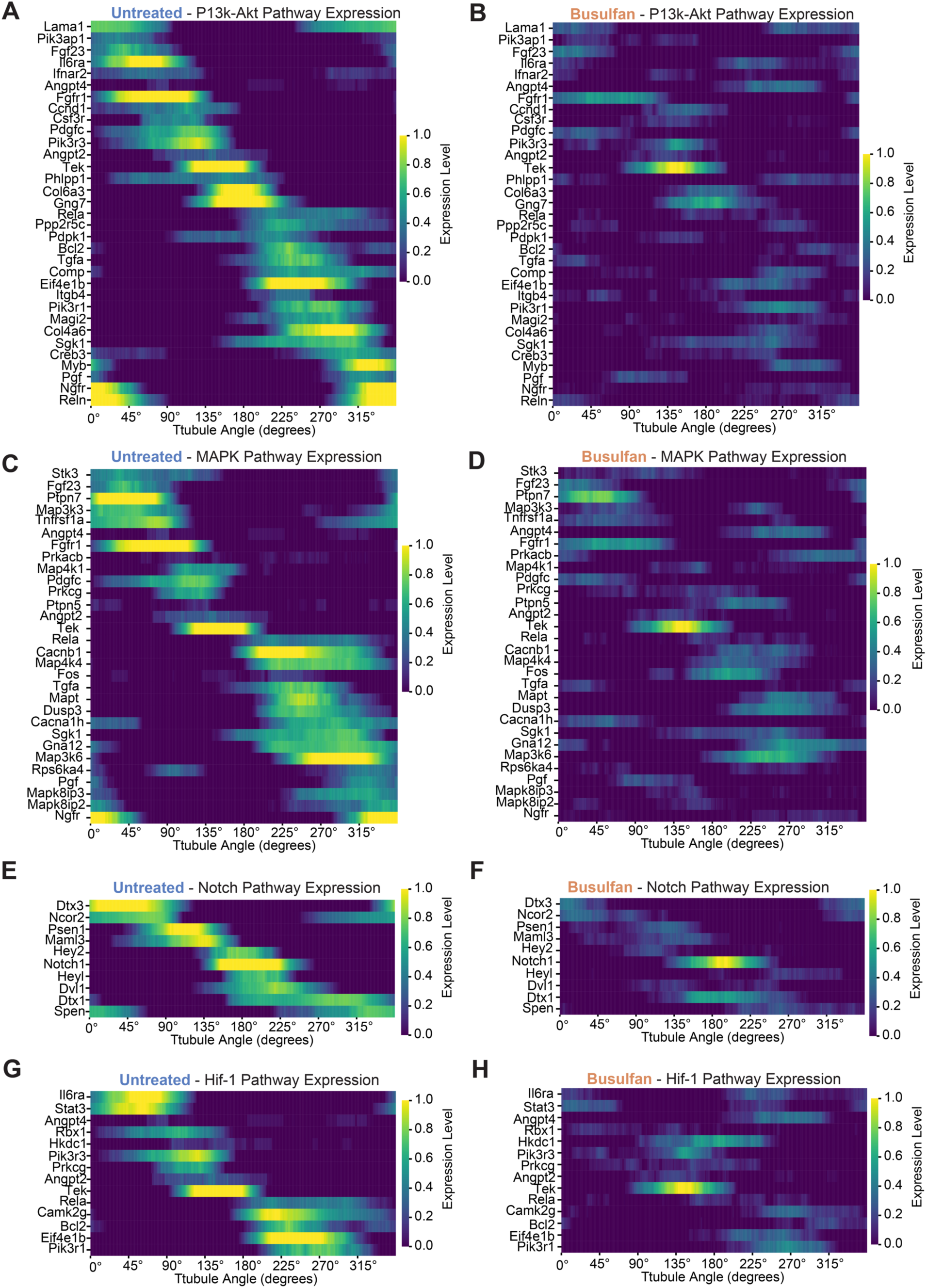
Gene expression across downregulated pathways between Sertoli cells in Untreated and Busulfan conditions. (A-H) Heatmaps showing expression for genes across pathways showing significant differences between untreated and busulfan conditions. Displayed genes amongst the top 200 genes with the most difference in expression patterns compared to untreated as computed using a Kolmogorov-Smirnov test.

**Figure S10:**
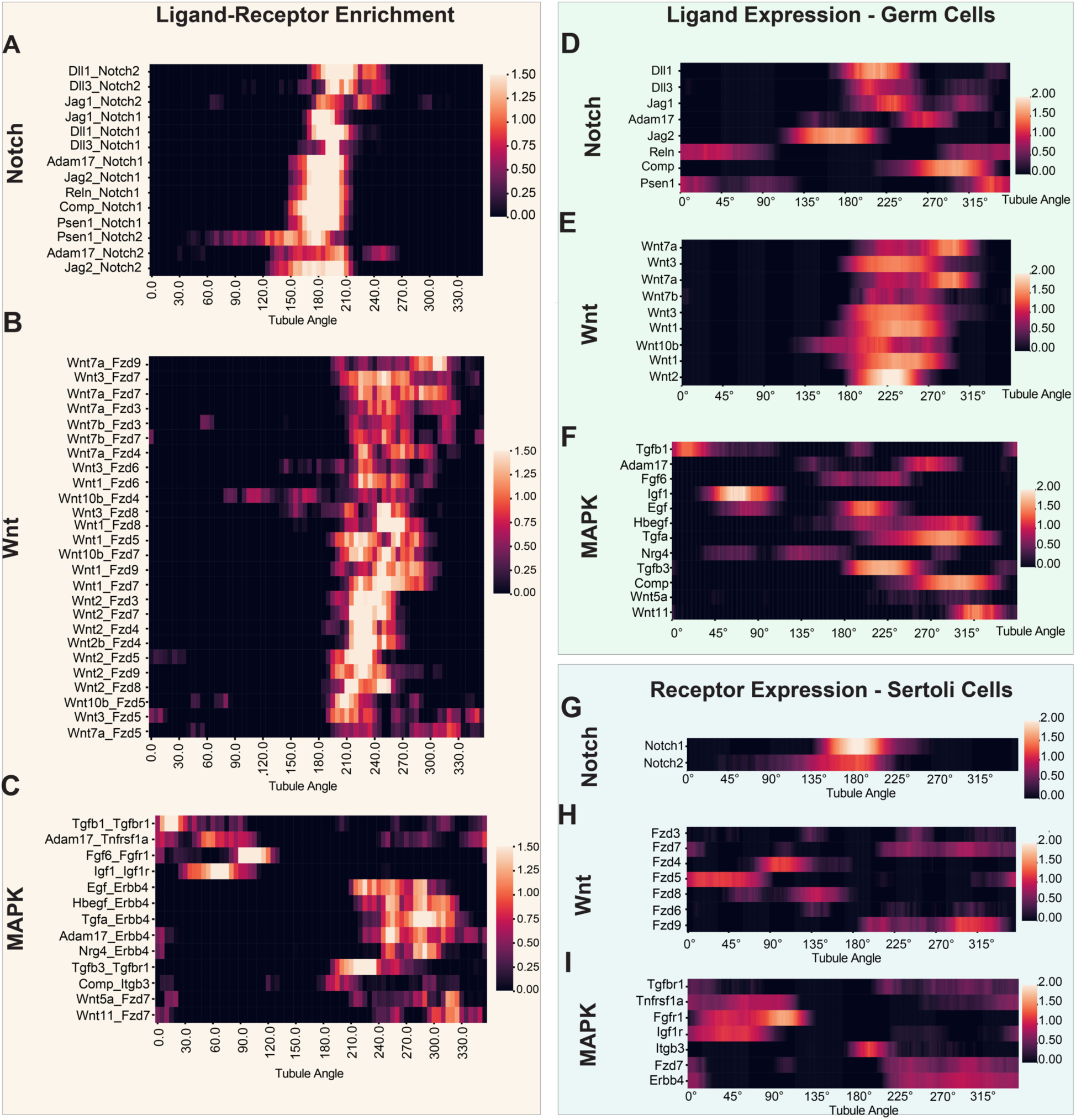
Ligand-Receptor interactions across Germ cells and Sertoli cells. (A -. **C)** Enriched Ligand-receptor interactions across Notch signaling pathway. (A) shows the aggregate signal of the interactions while (B) and (C) show the expression of ligands in germ cells and the receptors in Sertoli cells, respectively. **(D-F)** Enriched Ligand-receptor interactions across the Wnt signaling pathway. (A) shows the aggregate signal of the interactions while (B) and (C) the receptors in Sertoli cells show the expression of ligands in germ cells respectively. **(G - I)** Enriched Ligand-receptor interactions across the MAPK signaling pathway. (A) shows the aggregate signal of the interactions while (B) and (C) show the receptors in Sertoli cells show the expression of ligands in germ cells respectively.

**Figure S11:**
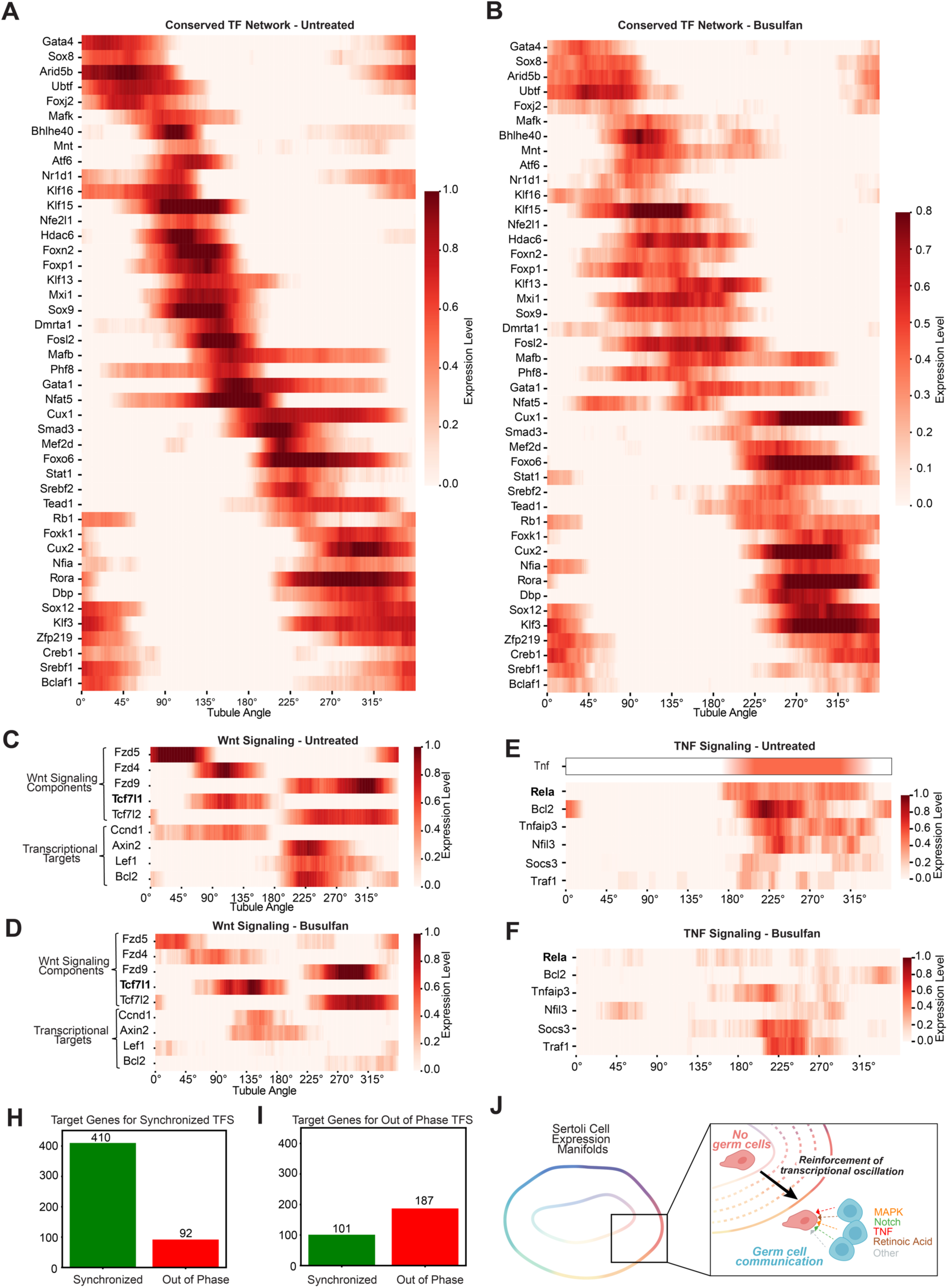
Conserved TF network and Extended Signaling Heatmaps. (A -. **B)** Heatmaps showing expression along Tubule Angle for TFs in conserved network(Fig 7G) in Sertoli cells in the untreated and busulfan condition. **(C - D)** Heatmaps showing expression along Tubule Angle for genes involved with Wnt Signaling in Sertoli cells in untreated and busulfan conditions. While Sertoli cells in the busulfan conditions still maintain signaling components including receptors and transcription factors (*Fzd5, Fzd4, Fzd9, Tcf7l1, Tcf7l2)*, downstream transcriptional targets are not expressed due to lack of signaling from germ cells (*Ccnd1, Axin2, Lef1, Bcl2).* **(E-F)** Heatmaps showing expression along Tubule Angle for genes involved with TNF Signaling in Sertoli cells in untreated and busulfan conditions. TNF is known to be expressed between stages VII - XI in germ cells^74^, and results in expression of canonical target genes *Rela* and *Bcl2.* Other genes *Tnfaip3, Nfil3, Socs3, Traf1* are all involved in the TNF signaling pathway. In Sertoli cells in the busulfan condition *Tnfaip3, Nfil3* show downregulation, while *Socs3 and Traf1* show upregulation. *Socs3,* in particular has been shown to be upregulated as a compensatory mechanism to downregulate cytokine signaling due to lack of TNF signaling^96^. **(H - I)** Histograms quantifying the number of target genes that were synchronized or out-of-phase between the Sertoli cells in the untreated and busulfan conditions. Synchronized was defined as having a Pearson correlation along tubule angle >0.80 and a peak angle that was within 40 degrees in both datasets. Out of phase was defined as fitting neither of these categories. **(J)** Schematic of overall model; Sertoli cells in the absence of germ cells maintain an innate cyclic transcriptional profile. With communication of germ cells, including signaling via MAPK, Notch, TNF, Wnt, retinoic acid, and other signaling mechanisms, germ cells can reinforce this cyclic transcriptional profile. At a topology level, the innate cyclic transcriptional profile occupies a space within the reinforced cyclic transcriptional profile in the presence of germ cells.

## Notes

### Summary of Updates

Added a few citations that were erroneously missing.

